# Neural categorization of visual words of alphabetic and non-alphabetic languages

**DOI:** 10.1101/2025.09.30.679494

**Authors:** Guo Zheng, Shihui Han

## Abstract

Languages provide social-category markers that tag people as one or another social group. How does the brain sort words into different language categories as a basis of the social-categorization function of language? We addressed this issue by testing neural categorization of visual words of different writing systems in nine studies using electroencephalography, magnetoencephalography, and a repetition suppression paradigm. We showed that a neural network, including the anterior temporal, insular, orbital frontal, and ventral occipito-temporal cortices in both hemispheres, was engaged in computations of correlation distances between two words to represent intra-language similarity and inter-language difference during categorization of visual words of alphabetic and non-alphabetic languages. These processes occurred as early as 150 ms post-stimulus, recruited within-hemisphere functional connections, operated independently of words’ semantic meanings and pronunciations, and exhibited consistently across individuals with diverse language backgrounds. These findings highlight the neural mechanisms of language-based spontaneous neural categorization of visual words as a basis of the social-categorization function of language.

## Introduction

Language is a sophisticated system for transmission of information (e.g., thoughts, knowledge, and feelings) among individuals (Fedorenko et al., 2024b). This linguistic communication function of language relies on the processing of semantic meanings and pronunciations of spoken or written words. Languages also provide social-category markers that tag individuals who use different languages as separate social groups (Bucholtz and Hall, 2010; DeJesus et al., 2018; Kinzler, 2021; Kinzler and Dautel, 2012; Kinzler et al., 2007; Pietraszewski and Schwartz, 2014; Roberts, 2013). This social-categorization function of language has notable consequences in human societies. For example, language serves as a key dimension of ethnic group identity which in turn influences social behaviors (Sachdev and Bourhis, 1990; Scherer and Giles, 1979). In extreme cases, individuals who spoke a different language were classified into an out-group and persecuted (Greenberg, 2008; Shell, 2001). The social-categorization function of language emerges early during human development (Kinzler, 2021; Rhodes and Baron, 2019). Infants expect two individuals who speak the same language to be affiliated (Liberman et al., 2017b) and favor native over non-native speakers during social learning (Howard et al., 2015) and acts of giving (Kinzler et al., 2012). Studies of adults also revealed evidence that language is used as a social cue for categorization of perceived faces (Baus et al., 2021; Champoux-Larsson et al., 2022) and social categorization of faces based on language may occur automatically (Lorenzoni et al., 2022). The social-categorization function of language revealed in these behavioral studies implicates that rapid categorization of words of different languages may occur in the human brain. Furthermore, the findings of infant studies (e. g., Liberman et al., 2017b) suggest that the neural process involved in categorization of words of different languages may develop even prior to the processing of linguistic properties (e.g. semantic meanings) of words.

Nevertheless, up to date, there has been little neuroimaging research examining the neural mechanisms underlying automatic and fast categorization of words of different languages.

Language-based categorization of written or spoken words with different pronunciations or semantic meanings lays the foundation for knowing who use the same or different languages and thus belong to the same or diverse social groups. The neural mechanisms of language-based categorization of words have been overlooked in previous research that usually focused on the linguistic communication function of language. Linguistic studies have revealed a core neural network that is dominated by the left hemisphere (including the inferior/middle frontal gyri and superior/middle temporal gyri) and enables the processing of linguistic properties (e.g., pronunciations and semantic meanings) of words (Fedorenko et al., 2024a; Friederici and Gierhan, 2013). This network responds to diverse languages (Malik-Moraleda et al., 2022; Siok et al., 2004), activates similarly to native and non-native languages (Li et al., 2021; Malik-Moraleda et al., 2024), and exhibits similar patterns of responses to spoken and written words/sentences (Fedorenko et al., 2024a). Electroencephalography (EEG) and magnetoencephalography (MEG) studies have revealed that visual words initially activate the left ventral occipito-temporal cortex at approximately 170 ms after stimulus onset, reflecting pre-lexical word form processing (Marinkovic et al., 2003; Nan et al., 2022; Pylkkänen and Marantz, 2003). The activation then spreads to the left superior temporal sulcus (LSTS)/inferolateral temporal area at ∼230 ms and anterior temporal lobe (ATL) at ∼350 ms, and then to the bilateral inferior prefrontal cortices and orbitofrontal cortices (OFC) at ∼400 ms (Marinkovic et al., 2003). The left temporal and frontal activities are particularly important for semantic and phonological processing of words and sentences (Hodgson et al., 2021) around 400 ms after word presentation (Zhu et al., 2022) as well as social-semantic working-memory (Zhang et al., 2023a). Because even words of an unlearned language can serve as a social-category marker of a ‘not-us group’, the neural dynamics of language-based categorization of words may be different from that involved in the processing of linguistic properties of words. The brain regions underlying social concept/knowledge and social categorization (Arioli et al., 2021; Olson et al., 2013; Pang et al., 2025; Pobric et al., 2016; Zahn et al., 2007; Zhou et al., 2020) may play important roles in categorization of words of different languages.

The present study investigated neural dynamics of categorization of visual words of two different (an alphabetic versus a non-alphabetic, or two different alphabetic) languages by combining EEG/MEG with a repetition suppression (RS) paradigm adopted from previous studies of social categorization of faces (Zhang et al., 2023b; Zhou et al., 2020). RS refers to the attenuation in neural responses to a repeated occurrence of stimuli that engage common neuronal populations or processes due to habituation. The RS paradigm consisted of an alternating condition (Alt-Cond), in which visual words of two different languages were presented alternately, and a repetition condition (Rep-Cond), in which words of one language were presented repeatedly (Fig. 1a). Neural responses to stimuli of the same category were attenuated in the Rep-Cond compared to Alt-Cond due to habituation and this RS effect has been examined to disentangle the neural activities underlying categorization of faces and body silhouettes of a specific social group (Pu and Han, 2025; Zhang and Han, 2021; Zhang et al., 2023b; Zhou et al., 2020).

**Figure 1.**
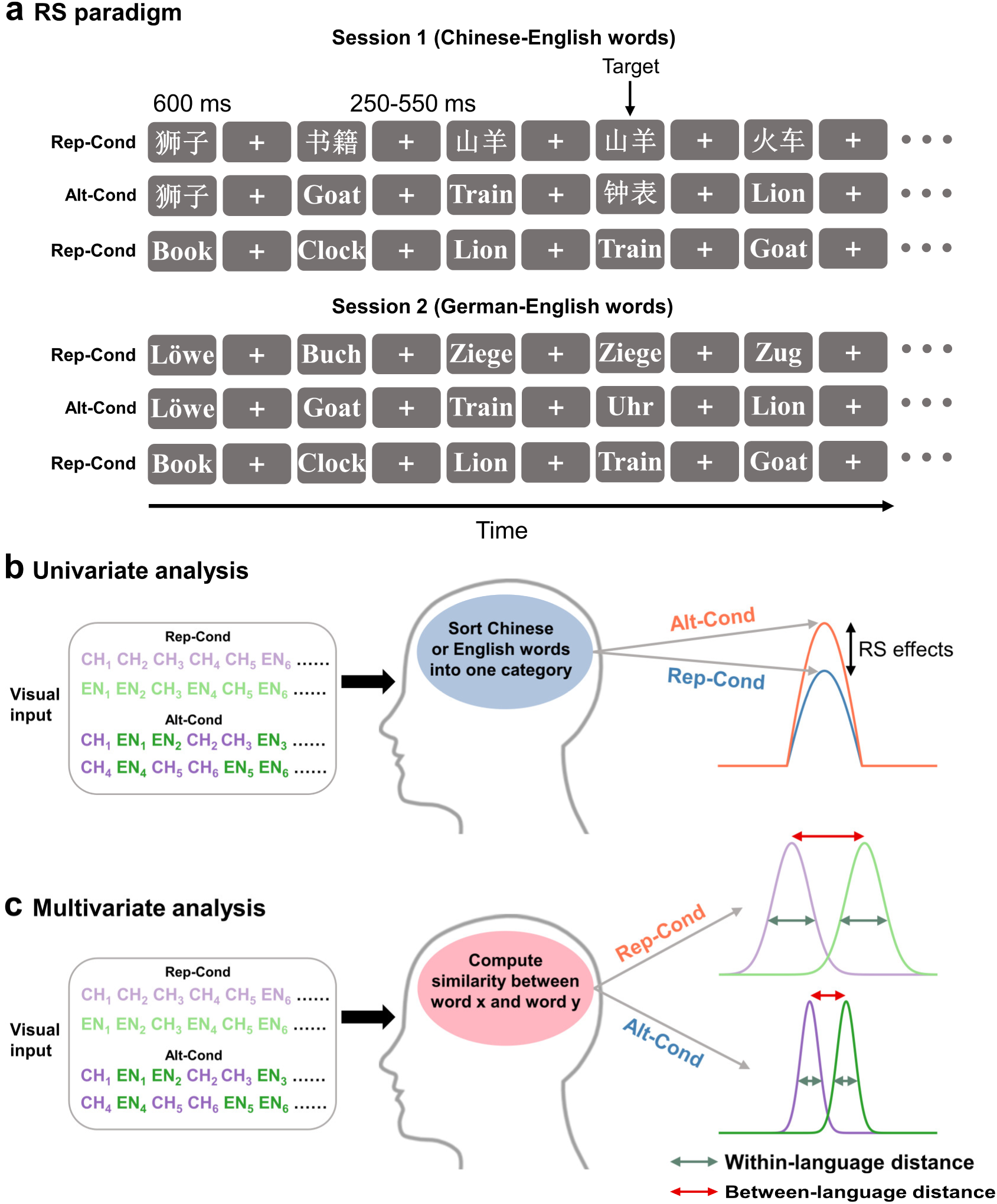
Illustrations of the RS paradigm and univariate/multivariate analyses of brain activities in response to words. (a) The RS paradigm. Words of two different languages (e.g., Chinese vs. English, or English vs. German) were presented alternately in the Alt-Cond, and words of one language were presented repeatedly in the Rep-Cond. (b) Univariate analyses. The RS effect was quantified as the decrease of averaged neural responses to words of one language in the Rep-Cond vs. Alt-Cond.

In nine experiments we recorded EEG/MEG signals from Chinese, English, and German speakers when viewing words of an alphabetic language and a non-alphabetic language (English and Chinese words, or Italian and Korean words) or of two alphabetic languages (English and German) in the Rep-Cond and Alt-Cond. We recorded EEG signals from Chinese participants to examine temporal neural dynamics of spontaneous language-based word categorization in Experiment 1. The similar paradigm was employed in Experiments 2 and 3 to investigate whether perceptual features or radical/letters of words are sufficient to generate spontaneous language-based categorization of visual words. The results in Experiment 1 were replicated in native English and German speakers in Experiments 4 and 5, respectively. Neural dynamics of categorization of words of two unlearned languages were further investigated in Chinese participants in Experiment 6. Finally, the neural networks supporting the spontaneous categorization of words of two learned or unlearned languages were localized using MEG in Chinese and English speakers in Experiments 7-9, respectively.

We performed univariate analyses of the RS effects on EEG and MEG signals in response to words of the same language to examine both timing and architecture of the integrated neural activities related to language-based word categorization (Fig. 1b). Since visual categorization of objects or faces depends on the processing of both within-category similarity (or intra-group) and between-category (or inter-group) difference between perceived stimuli (Freedman et al., 2001; Ito and Bartholow, 2009; Kriegeskorte et al., 2008; Zhou et al., 2020), we further conducted multivariate representational similarity analyses (Kriegeskorte et al., 2006) of correlation distances between two words to disentangle neural processes of intra-language similarity and inter-language difference between words that are fundamental to language-based word categorization. The processing of intra-language similarity occurs when two words of the same language are perceived repeatedly with short interstimulus intervals.

Because words of the same language were repeatedly presented in the Rep-Cond and words of two different languages were displayed in the Alt-Cond, the processing of intra-language similarity occurred more frequently and would be inhibited in the Rep-Cond (vs. Alt-Cond) due to habituation (Fig. 1c). By contrast, the processing of inter-language difference takes place when two words of different languages are perceived with short interstimulus intervals. Since words of different languages appeared more frequently in the Alt-Cond (vs. Rep-Cond), we would expect RS of the processing of inter-language difference in the Alt-Cond (vs. Rep-Cond). The neural processing of intra-language similarity was quantified as correlation distances between neural responses to two words of the same language whereas the neural processing of inter-language difference was assessed as correlation distances between neural responses to two words of two different languages. The correlation distances from the multivariate analyses were further employed to assess how words of one language are clustered and how far words of two languages are separated in a two-dimensional (2D) space during language-based word categorization. Enhanced language-based word categorization is associated with smaller intra-language correlation distances, which reflect more densely clustered words of the same language, and larger inter-language correlation distances, which manifest further separated words of two different languages. This approach is different from previous research that focused on distinct neural responses to visual words of two different languages in the brain regions such as the fusiform/lingual gyri that are specific to word-form processing (Zhan et al., 2023).

This RS effect manifested habituation of the integrated neural activities that supported classification words into one or another category and occurred more frequently in the Rep-Cond (vs. Alt-Cond). (c) Multivariate analyses. Correlation distances between neural responses to two words were calculated to estimate how words of one language were clustered (i.e., intra-language similarity) and how words of two languages were separated (i.e., inter-language difference) during language-based word categorization.

## Results

### Temporal dynamics of spontaneous language-based word categorization

To examine temporal characteristics of the neural processes involved in spontaneous language-based word categorization, in Experiment 1, we recorded EEG in two sessions from native Chinese speakers who learned English but not German (N=34, see Table S1 for information about all participants in our work). In Session 1 Chinese words (animal and tool names) and English words (translated from the Chinese words, see Table S2 for all the stimuli used in our study) were presented in the Alt-Cond and Rep-Cond, respectively (Fig. 1a). The same design was applied to English words and German words (translated from the Chinese words) in Session 2. Participants responded to a casual target word that was presented repeatedly in two consecutive trials by pressing a button during EEG recording. This one-back task required neither processing of linguistic properties nor intentional classification of words and thus allowed us to examine neural activities underlying spontaneous language-based word categorization. Behavioral performances in the one-back task did not differ significantly between words of different languages (see Table S3 for details of behavioral results in all studies), indicating comparable attentional demand and task difficulty in detection of target words of different languages.

Event-related brain potentials (ERPs) in response to non-target words were characterized by an early negative activity at 90–140 ms (N1), a positive activity at 140–280 ms (P2), and a late negative activity at 280–350 ms (N2) over the frontal/central electrodes, a long-latency positivity at 400–600 ms (LPP) at the centro-parietal electrodes, and a negative activity at 150–210 ms (N170) at the occipito-temporal electrodes (Fig. 2a and 2b, Fig. S1). We conducted univariate whole-brain cluster-based permutation *t*-tests to examine the RS effect on neural responses to non-target words (i.e., decreased amplitudes in the Rep-Cond vs. Alt-Cond). The results in Session 1 revealed significant clusters showing the RS effects over the middle frontal/central/parietal regions at 134–320 ms for Chinese words and at 172–300 ms for English words (a pre-defined threshold of *P* < 0.05, cluster-level *P* < 0.05, two-tailed, 10,000 iterations). However, similar analyses of brain activities in Session 2 did not show reliable RS of neural responses to English and German words (Fig. S2). These results provided electrophysiological evidence for spontaneous language-based categorization of words as early as 150 ms after stimulus onset between an alphabetic language and a nonalphabetic language (i.e., English and Chinese) but not between two alphabetic languages (i.e., English and German).

**Figure 2.**
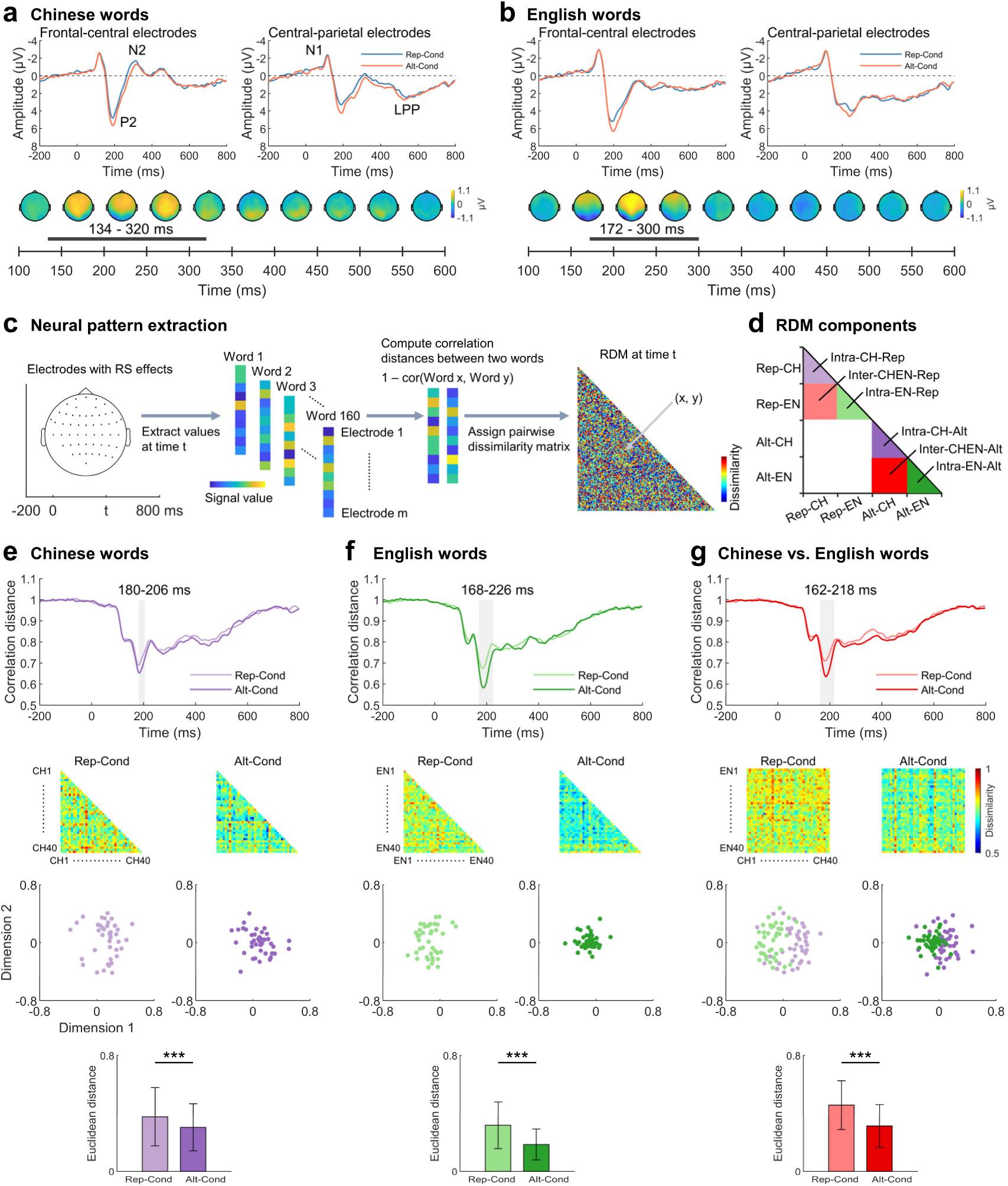
EEG results of Chinese speakers in Experiment 1. (a) and (b) Results of univariate analyses. Top panels illustrate electrophysiological responses to Chinese and English words in the Alt-Cond and Rep-Cond, respectively. Bottom panels show scalp distributions of significant RS effects on neural responses to words of each language. (c) Illustration of the procedure of computing neural RDMs in the multivariate analyses of correlation distances between words. (d) Illustration of the 160 × 160 neural RDM. Triangles represent the neural RDMs corresponding to intra-language similarity. Squares represent the RDMs corresponding to inter-language difference. (e), (f), and (g) Results of multivariate and multidimensional scaling analyses. The top two panels show the time courses of significant differences in correlation distances between words corresponding to intra-language similarity and inter-language difference between the Alt-Cond and Rep-Cond and the neural RDM in the two conditions, respectively. The bottom two panels illustrate clustered representations of the words in the 2D word space built based on the first two dimensions of multidimensional scaling analyses of neural RDMs corresponding to intra-language similarity and inter-language difference, respectively, and the mean Euclidean distances in the 2D word space between two words of the same language and between two words of different languages. *** *P* < 0.001.

Next, we conducted multivariate representational similarity analyses of the neural responses to words to examine the neural processes of intra-language similarity and inter-language difference between words, respectively. A 160 × 160 neural representation dissimilarity matrix (RDM), including 40 words from each language in the Alt-Cond and Rep-Cond, was constructed using the EEG data at the electrodes at which the ERP amplitudes showed the significant RS effects in the univariate analyses (Fig. 2c and 2d). This RDM consisted of a matrix of correlation distances (calculated as 1–Pearson correlation coefficients) between neural responses to two words of the same language, which represents the neural response pattern of the processing of intra-language similarity (a smaller intra-language correlation distance corresponds to a more clustered representation of words of the same language). The RDM also had a matrix of correlation distances between neural responses to two words of different languages, which represents a neural response pattern of the processing of inter-language difference (a larger inter-language correlation distance corresponds to greater separation of representations of words of two languages). As predicted, permutation *t*-tests showed that the mean correlation distance of the RDMs corresponding to intra-language similarity was significantly increased in the Rep-Cond (vs. Alt-Cond) at 180–206 ms for Chinese words and at 168–226 ms for English words (Fig. 2e-g, see Fig. S3 for these RS effects at the individual level), indicating weakened clustered representations of Chinese (or English) words in the Rep-Cond (vs. Alt-Cond) due to habituation. By contrast, the mean correlation distance of the RDMs corresponding to inter-language difference was significantly reduced in the Alt-Cond (vs. Rep-Cond) at 162–218 ms, indicating more closed representations of Chinese and English words in the Alt-Cond (vs. Rep-Cond) due to habituation.

Because these neural RS effects were consistently observed within 300 ms, the following multivariate analyses focused on the results in this time window.

To further assess the neural categorical representations of Chinese and English words in the Alt-Cond and Rep-Cond, we conducted multidimensional scaling analyses of the 160 × 160 RDMs averaged in the time window of the significant RS effects. The first two components of the results of these analyses were then used to construct 2D word spaces in which words of the same language or of the two different languages were plotted. As can be seen in Fig. 2e-g, words of the same language are clustered more densely in the Alt-Cond (vs. Rep-Cond) whereas words of the two different languages separate more distantly in the Rep-Cond (vs. Alt-Cond). These differences were further quantified by comparing the mean Euclidean distances in the word spaces between two words of the same language and between two words of different languages in the Alt-Cond and Rep-Cond. Together, the results of our multivariate analyses established two neural processes of intra-language similarity and inter-language difference that took place spontaneously during categorization of Chinese and English words in Chinese speakers.

### Perceptual features or radical/letters of words are not sufficient to generate spontaneous language-based categorization of words

Did the neural RS effects observed in Experiment 1 manifest habituation of the processing of non-specific perceptual shape features of words? We clarified this issue in Experiment 2 by creating two sets of scrambled stimuli from the Chinese and English words used in Experiment 1. The scrambled stimuli possess both global and local shape features of the words but lack word-specific information (e.g., semantic meanings and language categories). We recorded EEG signals in response to the scrambled stimuli from an independent sample of Chinese speakers (N=34) using the same experimental procedure as that in Experiment 1. Whole-brain cluster-based permutation *t*-tests did not find any significant RS effect on neural responses to the scrambled stimuli (Fig. S4), indicating that perceptual features of visual words contribute little to the neural RS effects related to spontaneous categorization of Chinese and English words.

In Experiment 3 we further tested whether categorization of radicals of Chinese words and letters of English words engages similar neural processes as those involved in categorization of Chinese and English words. We recorded EEG signals in response to Chinese radicals and English letters from an independent sample of Chinese speakers (N=34) using the same experimental procedure as that in Experiment 1.

Univariate whole brain cluster-based permutation *t*-tests showed significant RS effects on the ERP amplitudes to Chinese radicals at 172–286 ms over the central region and to English letters at 214–364 ms over the occipital regions (Fig. S5a and S5b).

Multivariate analyses of the correlation distances corresponding to intra-radical similarity did not show any significant difference between the Rep-Cond and Alt-Cond. The mean correlation distances corresponding to intra-letter similarity and inter-radical-letter difference were significantly decreased in the Rep-Cond than Alt-Cond but in time windows delayed (after 230 ms) compared to those observed for Chinese and English words in Experiment 1 (Fig. S5c-e). These results provide no evidence that perception of the middle-level units of Chinese and English words (i.e., radicals and letters) employs the same early fronto-central neural processes as those involved in spontaneous categorization of Chinese and English words.

### Neural dynamics of language-based word categorization is independent of people’s language backgrounds

To generalize the findings in Experiment 1 to populations of different language backgrounds, we recorded EEG signals to words from native English speakers (Experiment 4, N=34) who learned Chinese (but not German) and native German speakers (Experiment 5, N=34) who learned both Chinese and English. The stimuli and procedure were the same as those in Experiment 1. The results of both univariate and multivariate analyses replicated those observed in Experiment 1 (see Fig. S6–S9 for details), indicating similar neural processes involved in language-based words categorization regardless of speakers’ language proficiency and learning experiences.

### Neural dynamics of categorization of words of two unlearned languages

So far, the neural RS effects in Experiments 1, 4, and 5 were observed for words of two learned languages. In Experiment 6 we further investigated to what degree semantic meanings and pronunciations of words contributed to the neural processes of intra-language similarity and inter-language difference during spontaneous language-based word categorization. We recorded EEG signals in response to Korean and Italian words (see Table S2) from an independent sample of Chinese speakers (N=34) who had not learned Korean and Italian when being tested. The experimental procedure was the same as that in Experiment 1. The participants were informed of viewing Korean and Italian words during EEG recording but did not know semantic meanings of these words and were unable to pronounce the Korean words (though might be able to pronounce Italian words in the way to spell English words since they had learned English). If semantic meanings or pronunciations of words are necessary for spontaneous language-based categorization of words, the neural RS effects would not occur to Korean or Italian words.

Cluster-based permutation *t*-tests of the ERP amplitudes to non-target revealed significant RS effects over the middle frontal/central/parietal regions at 136–310 ms for Korean words and at 156–514 ms for Italian non-target words (Fig. S10a and S10b). Multivariate analyses of neural responses to words showed a significantly increased mean correlation distance corresponding to intra-language similarity in the Rep-Cond (vs. Alt-Cond) at 164–216 ms for Korean words and at 166–246 ms for Italian words (Fig. S10c-e). Moreover, the mean correlation distance corresponding to inter-language difference was significantly reduced in the Alt-Cond (vs. Rep-Cond) at 164–236 ms. Similarly, the multidimensional scaling analyses of the RDMs revealed more densely clustered representations of words of the same language in the Alt-Cond (vs. Rep-Cond) and more distantly separated representations of words of the two different languages in the Rep-Cond (vs. Alt-Cond) in the 2D word space. These results demonstrated that spontaneous language-based categorization also occurred to words of two unlearned languages. Furthermore, the time courses of neural processes of intra-language similarity and inter-language difference involved in categorization of Korean and Italian words were akin to those of categorization of words of two learned languages (i.e., Chinese and English). These results indicate that semantic meanings or pronunciations of words contribute little to spontaneous categorization of words of an alphabetic language and a non-alphabetic language.

### A neural network underlying language-based categorization of words

Next, we sought to localize the neural network underlying spontaneous language-based categorization of words in Experiments 7a and 8a. We recorded 306-channel, whole-head anatomically constrained MEG signals in response to Chinese and English words from two independent samples of native Chinese and English speakers (N=34 in each group). High-resolution structural MRI was combined with temporally precise whole-head high-density MEG to localize brain regions in which activities showed RS effects and to examine the functional roles of this network in processing intra-language similarity and inter-language difference between words. The stimuli and procedures were the same as those used in Experiment 1. Because our EEG data analyses showed consistent neural RS effects in Chinese and English speakers, we combined Chinese and English speakers’ MEG data to examine the neural RS effects. We first assessed time courses of the RS effect on sensor-space MEG signals to words by pooling across Chinese and English words. A whole-brain cluster-based permutation *t*-test of sensor-space MEG signals in the Rep-Cond and Alt-Cond revealed three significant clusters (magnetometer signals:136–278 ms and 151–268 ms; gradiometer signals: 154–257 ms, using a predefined threshold of *P* < 0.01, two-tailed, 10,000 iterations, and a cluster-level threshold of *P* < 0.05, Fig. 3a). These results provide MEG replication of our EEG findings of the temporal dynamics of spontaneous language-based word categorization.

**Figure 3.**
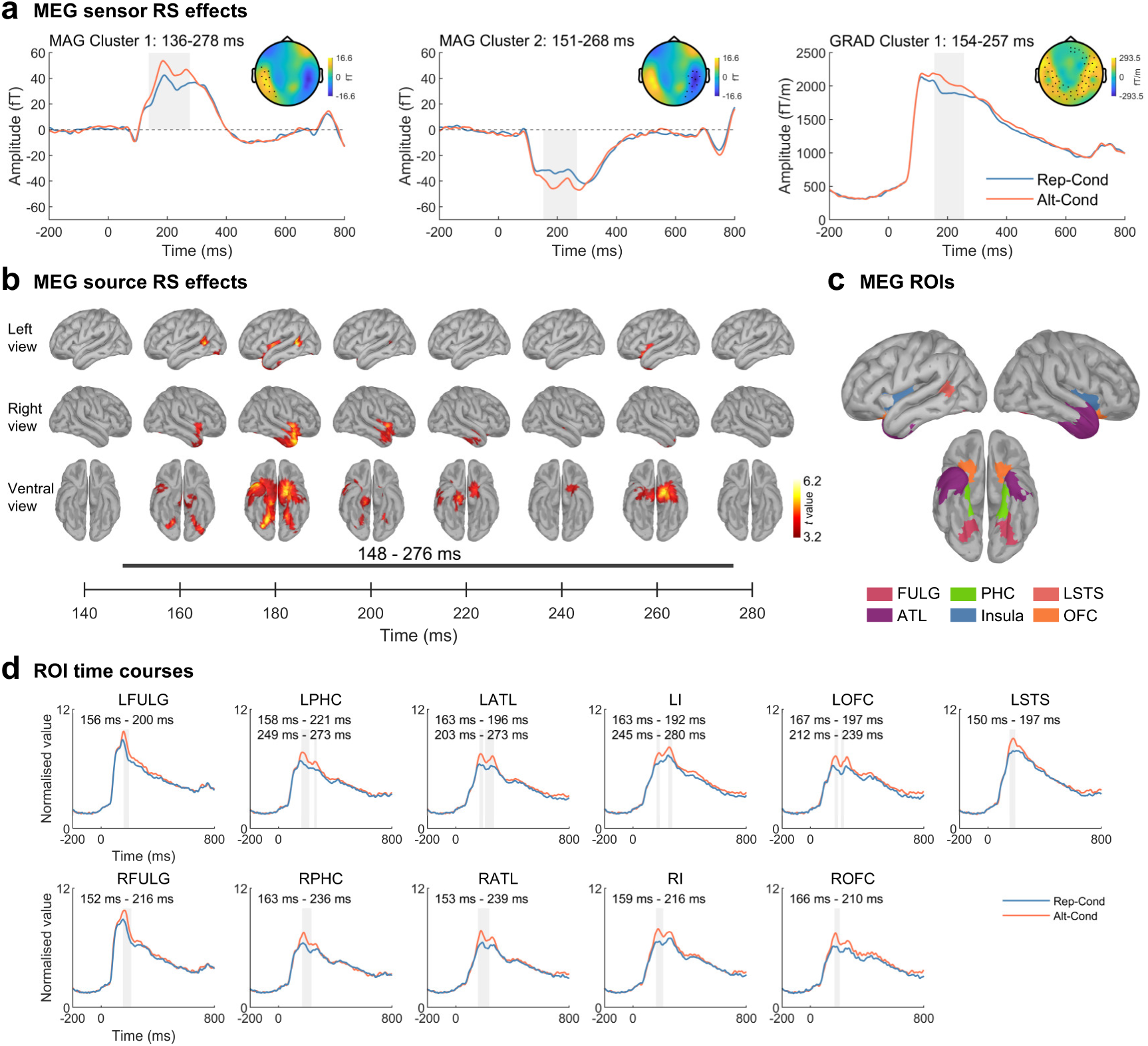
MEG results combined Chinese and English words across Chinese and English speakers in Experiments 7a and 8a. (a) Significant clusters of RS effects on magnetometer signals (the left two panels) and gradiometer signals (the right panel). (b) Results of the whole-brain source analyses. Shown are left, right, and ventral views of brain regions in which MEG signals in response to words of the same language decreased significantly in the Rep-Cond (vs. Alt-Cond). (c) Illustration of the regions of interest used in the following MEG data analyses. (d) Source-space MEG signals in the brain regions showing significant RS effects. Time windows of the significant clusters are shown for each brain region. LFULG=left fusiform and lingual gyrus; RFULG= right fusiform and lingual gyrus; LPHC=left parahippocampal cortex; RPHC=right parahippocampal cortex; LATL=left anterior temporal lobe; RATL=right anterior temporal lobe; LI=left insula; RI=right insula; LOFC=left orbital frontal cortex; ROFC=right orbital frontal cortex; LSTS=left superior temporal sulcus;

To probe the neural network involved in language-based categorization of words, we performed source reconstruction of MEG signals and then combined source-space MEG signals to Chinese and English words across Chinese and English speakers to search for brain regions in which responses to words of the same language were suppressed in the Rep-Cond (vs. Alt-Cond). The results of this univariate analysis revealed significant RS effects on activities in the bilateral ATLs, insula, OFC, occipito-temporal cortices, and the LSTS at a predefined threshold of *P* < 0.001, 10,000 iterations, and a cluster-level threshold of *P* < 0.05, one-tailed (Fig. 3b). To further assess the time courses and hemispheric asymmetry of the neural RS effects in different nodes of this network, we defined eleven regions of interest (ROIs) based on the intersection of brain regions showing significant RS effects and the Desikan-Killiany-Tourville atlas (Klein and Tourville, 2012), including the ATL, insula, OFC, fusiform and lingual gyrus (FULG), parahippocampal cortex (PHC) in both hemispheres, and LSTS (Fig. 3c). Cluster-based permutation *t*-tests of the activities in these brain regions within 300 ms after word onsets identified significant RS effects at 150–280 ms (Fig. 3d). The neural RS effects started slightly earlier in the right than left hemispheres (except the PHC), though a repeated measures analysis of variance (ANOVA) of the peak latency of the RS effect with Hemisphere (left vs. right) and Brain regions (OFC, insula, ATL, FULG, PHC) as independent within-subjects factors did not find a significant difference between the two hemispheres (*P* > 0.9). We also compared the magnitude of the RS effects in the two hemispheres by conducting ANOVA of the peak RS effects with Hemisphere (left vs. right) and Brain regions (OFC, insula, ATL, FULG, PHC) as independent within-subjects factors but failed to find a significant difference in the neural RS effects between the two hemispheres (*P* > 0.4). These results provide no evidence for dominance of one over the other hemisphere during language-based word categorization.

To further explore the functional role of this network in processing of intra-language similarity and inter-language difference between words, we constructed global network RDMs based on the correlation distances calculated from 313 subregions (each subregion had 5 vertices on average) from the eleven ROIs shown in Fig. 5c (using the mean source-space MEG signals in each subregion, see Methods for details, Fig. 4a). If this neural network serves the processing of intra-language similarity during language-based categorization of words, the network RDMs corresponding to intra-language similarity would be attenuated in the Rep-Cond (vs. Alt-Cond) whereas the network RDMs corresponding to inter-language difference would show a reverse pattern due to habituation. We tested these predictions by performing permutation *t*-tests of the time courses of the RDMs. The results revealed that the correlation distance corresponding to intra-language similarity was significantly increased in the Rep-Cond (vs. Alt-Cond) at 159–200 ms for Chinese words and at 151–198 ms for English words. Moreover, the correlation distance corresponding to inter-language difference was significantly reduced in the Alt-Cond (vs. Rep-Cond) at 150–232 ms and 236–294 ms (Fig. 4b-d, see Fig. S11 for these RS effects at the individual level). The multidimensional scaling analyses of the network RDM in the time windows of these significant clusters further unraveled more densely clustered representations of words of the same language in the Alt-Cond (vs. Rep-Cond) and more distantly separated representations of words of the two different languages in the Rep-Cond (vs. Alt-Cond) in the 2D word space. Similar results were obtained in separate analyses of MEG data in Experiments 7a and 8a, respectively (see Fig. S12 and S13). These results identified the word-categorization network that supported the processing of both intra-language similarity and inter-language difference during categorization of Chinese and English words.

**Figure 4.**
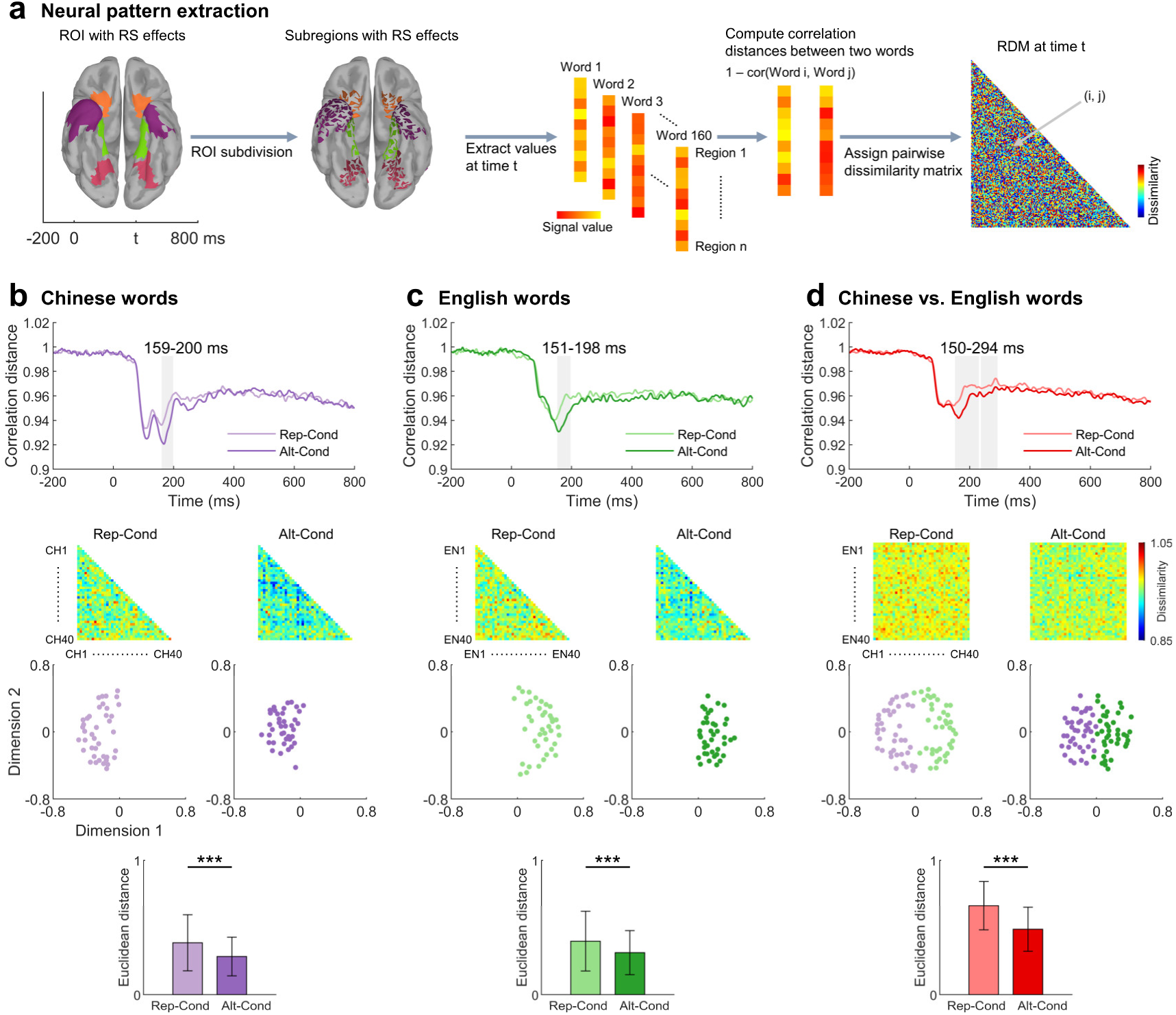
Results of the analyses of the global network RDM across Chinese and English speakers in Experiments 7a and 8a. (a) Illustration of the procedure to calculate network RDMs based on source-space MEG signals. (b), (c), and (d) Results of multivariate analyses. The top two panels show the time courses of significant differences in the correlation distances corresponding to intra-language similarity and inter-language difference between the Alt-Cond and Rep-Cond and the neural RDMs in the two conditions, respectively. The bottom two panels illustrate clustered representations of words in the 2D word space built based on the first two dimensions of multidimensional scaling analyses of neural RDMs corresponding to intra-language similarity and inter-language difference, respectively, and the mean Euclidean distances in the 2D word space between two words of the same language and between two words of different languages. \*\*\**P* < 0.001.

**Figure 5.**
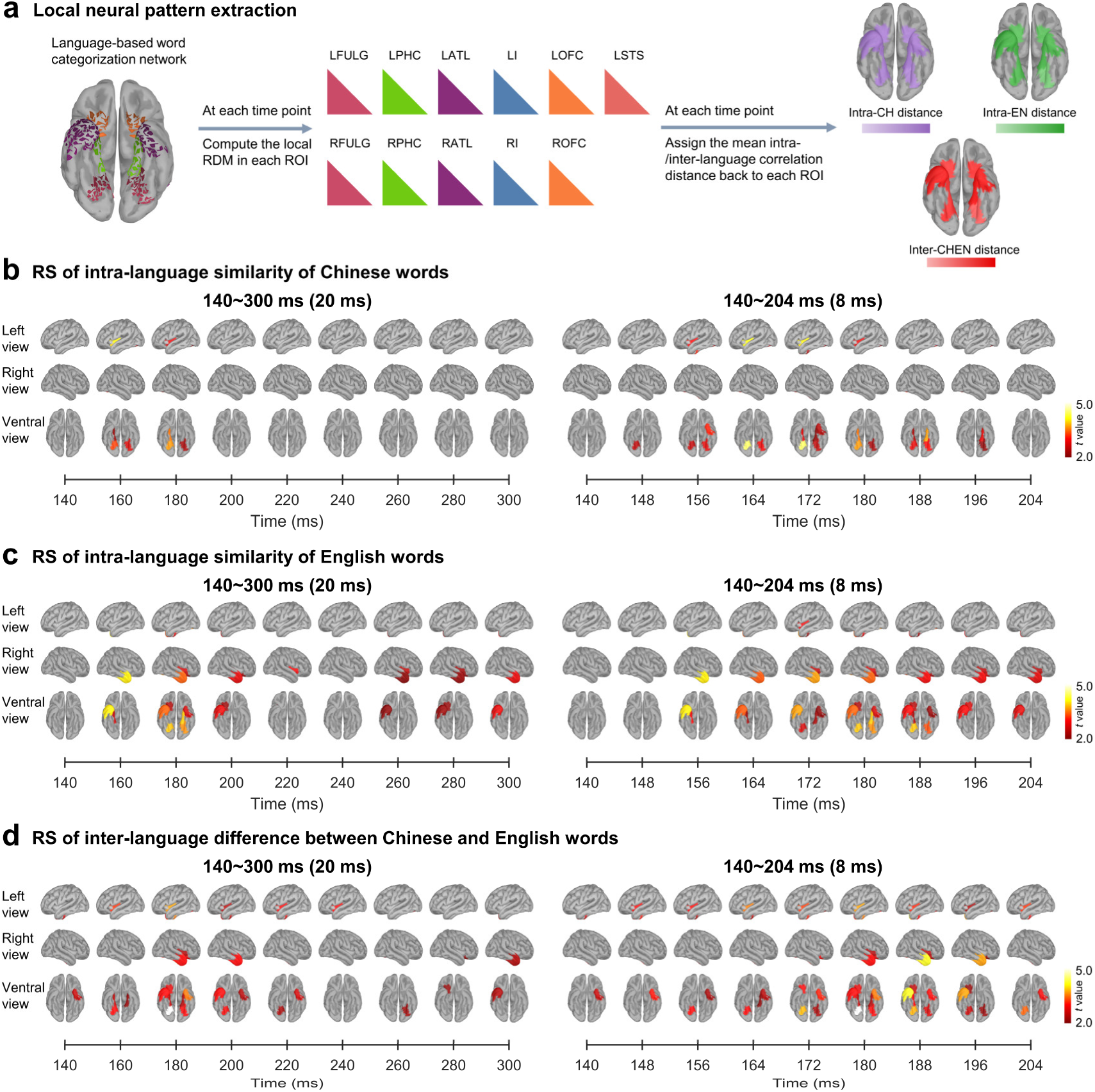
Results of the analyses of local RDMs across Chinese and English speakers in Experiments 7a and 8a. (a) Illustration of the procedure to calculate local RDMs based on source-space MEG signals. (b) and (c) Dynamic RS effects related to intra-language similarity of Chinese and English words in different ROIs. (d) Dynamic RS effects related to inter-language difference in different ROIs.

We further assessed dynamic contributions of each brain region to the processing of intra-language similarity and inter-language difference during language-based word categorization. We computed local RDMs using the patterns of activity in each of the 11 ROIs, respectively (see Methods for details). The mean correlation distance values specific to intra-language similarity and inter-language difference were calculated and assigned to all vertices in each ROI, as illustrated in Fig. 5a. We then performed whole-network cluster-based permutation *t*-tests to examine the dynamic RS effects on the mean correlation distance values (a predefined threshold of *P* < 0.025, 10,000 iterations, and a cluster-level threshold of *P* < 0.05, one-tailed). The results showed that the RS effects related to intra-language similarity initiated at 148 ms in the FULG for Chinese words and at 156 ms in the RATL for English words, and these early RS effects spread to other brain regions in the network (Fig. 5b and 5c). The RS effects pertaining to inter-language difference started at 140 ms in the LATL and then spread to other brain regions in the network (Fig. 5d). These results suggested distinct patterns of neural dynamics linked to the computations of intra-language similarity and inter-language difference.

We further performed a Granger causality analysis (GCA) (Barnett and Seth, 2014) to examine functional connectivity characteristics of the word-categorization network during the processing of intra-language similarity and inter-language difference between words. To this end, we calculated the time series of the RS effects on correlation distances between two words during 100–300 ms in each cell of the network RDM corresponding to intra-language similarity or inter-language difference in the eleven ROIs (Fig. 4c). These time series were then subject to GCA to estimate information flow among the ROIs during language-based word categorization (see Methods for details). The results revealed three connectivity characteristics of the network (Fig. 6a-c). First, functional connections occurred dominantly within the same hemisphere and no cross-hemisphere functional connection passed the same threshold. Second, forward and backward connections existed between adjacent brain regions (e.g., FULG and PHC, or ATL and OFC) whereas long-distance connections were rare. Third, while similar patterns of functional connections were observed for the processing of intra-language similarity and inter-language difference between words, the connectivity strength in the whole network was slightly but significantly greater during processing of intra-language similarity relative to inter-language difference. Finally, the connections were significantly stronger in the left than right hemispheres in connections from FULG to PHC, ATL to PHC, insula to ATL, and OFC to ATL, whereas the connection from ATL to insula was significantly stronger in the right than left hemispheres (*Ps* < 0.05, false discovery rate (FDR) corrected).

**Figure 6.**
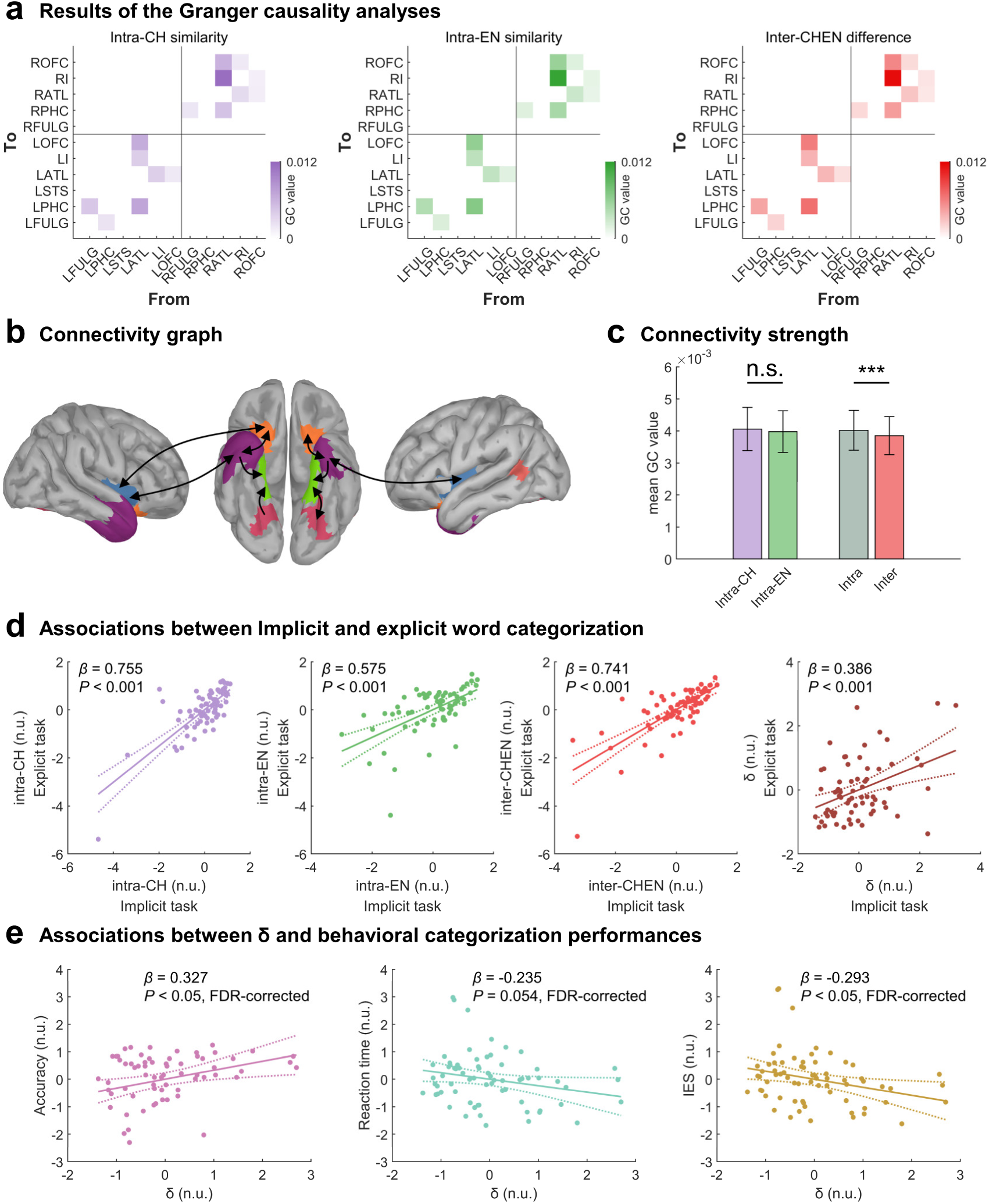
Results of GCA and linear regression analyses across Chinese and English speakers in Experiments 7a and 8a. (a) Connectivity values above a threshold (GC values > 0.001) corresponding to intra-language similarity and inter-language difference. Each square represents connectivity from one brain region indicated by the x-axis to another region indicated by the y-axis. (b) Patterns of functional connectivity between two ROIs. Arrows indicate directions of information flow from one to another brain region. Lines with two arrows indicate information flow with mutual directions. (c) The mean connectivity strength in the whole word-categorization network related to the processing of intra-language similarity and inter-language difference. (d) Associations between the correlation distances corresponding to inter-language difference and intra-language similarity, and the neural index of language-based word categorization (**δ**) during the implicit and explicit language-based word categorization tasks. Shown are partial regression plots (e) Associations between the neural index of language-based word categorization (**δ**) and behavioral performances (response accuracies, reaction time and inverse efficiency scores (IES)) during the explicit language-based word categorization tasks. *** *P* < 0.001; n.s. = no significance.

To further test the functional roles of neural computations of intra-language similarity and inter-language difference between words in language-based word categorization, we asked the participants in Experiments 7a and 8a to perform an explicit categorization task that required behavioral classification of Chinese and English words (Experiments 7b and 8b, see Methods for details). The participants sorted Chinese and English words with high accuracies and fast responses by pressing one of two buttons (Experiments 7b and 8b: accuracy = 96.04 ± 0.03% and 96.20 ± 0.03%, reaction time = 521 ± 64.3 and 481 ± 45.3 ms). If neural computations of intra-language similarity and inter-language difference are conducted spontaneously during both implicit and explicit language-based word categorization, intra-language similarity and inter-language difference computed in the Alt-Cond in an implicit task (i.e., the one-back task) would predict those computed in the explicit categorization task. In addition, participants with better behavioral classification performances in the explicit categorization task would be associated with smaller intra-language similarity (i.e., stronger clustered neural representations of words of one languages) but a larger inter-language difference (i.e., larger separation of neural representations of words of two languages).

To test these predictions, we extracted the neural activities from the word-categorization network identified in Experiments 7a and 8a during the explicit word categorization task in Experiments 7b and 8b. The correlation distances corresponding to inter-language difference and intra-language similarity were then calculated based on these activities in the time windows showing significant RS effects in the multivariate analyses in Experiments 7a and 8a. We then calculated a neural index (**δ**) that combined the contributions of inter-language difference and intra-language similarity to language-based word categorization:

**δ** = Inter_LD – (Intra_LS_CH_ + Intra_LS_EN_)/2 in which Inter_LD = inter-language difference; Intra_LS_CH_ = intra-language similarity of Chinese words; Intra_LS_EN_ = intra-language similarity of English words. A larger **δ** indicates more clustered neural representations of words of the same language and more separated neural representations of words of different languages. Behavioral performances during the explicit word categorization task were quantified by the mean response accuracy, mean reaction time, and the inverse efficiency score (IES, i.e., the ratio of the mean reaction time to response accuracy) that considered both the response accuracy and reaction time.

Linear regression analyses including Group (Chinese or English speakers) as a covariate showed that the correlation distances corresponding to intra-language similarity, inter-language difference and **δ** in the Alt-Cond in the one-back task in Experiments 7a and 8a positively predicted those in the explicit word categorization task in Experiments 7b and 8b (*Ps* < 0.001, Fig. 6d). These results suggest similar neural computations of intra-language similarity and inter-language difference in the same neural network during implicit and explicit language-based word categorization. More importantly, a larger **δ** calculated using the neural network responses to words in the explicit language-based word categorization significantly and positively predicted better behavioral categorization performances (i.e., higher accuracies: *P* < 0.05; shorter reaction times: *P* = 0.054, and smaller IES: *P* < 0.05, FDR-corrected, Fig. 6e), in Experiments 7b and 8b. However, the **δ** calculated using the source-space MEG signals in each of the eleven ROIs of the word-categorization network failed to predict behavioral performances during explicit language-based word categorization. These results indicate that the integrated computations of intra-language similarity and inter-language difference supported by the whole word-categorization network (rather than by a single region of the word-categorization network) provide a neural mechanism of behavioral categorization of words of an alphabetic language and a non-alphabetic language.

We also conducted a disruption analysis to examine which node of the word-categorization network was necessary for predicting behavioral categorization of words of two languages. To this end, we computed the **δ** of the word-categorization network by removing the bilateral FULG, PHC, ATL, insula, OFC, and LSTS, respectively. Thereafter, we conducted linear regression analyses to test whether **δ** of the disrupted network predicted behavioral categorization of words. The results showed that the association between **δ** and behavioral categorization of words was significant except when the bilateral FULGs were removed from the network (see Fig. S14), suggesting that the FULG may be more critical than the other nodes of the network in supporting behavioral performances during language-based categorization of words.

### The word-categorization network functions independently of words’ linguistic properties

Finally, we tested whether the neural network identified in Experiments 7a and 8a also serves language-based categorization of words of two unlearned languages in Experiment 9. We recorded MEG signals from an independent sample of Chinese speakers (N=34) who had not learned Korean and Italian when being tested. The stimuli and procedure were the same as those in Experiment 7 except that Korean and Italian words were used. The results in Experiment 9 replicated the results in Experiments 7a and 8a (Fig. S15–S17). These results provided further evidence that the word-categorization neural network appears to support the processing of intra-language similarity and inter-language difference during spontaneous categorization of words of an alphabetic language and a non-alphabetic language independently of the processing of words’ linguistic features (e.g., pronunciations or semantic meanings).

## Discussion

Together, our EEG/MEG findings uncovered the dynamic neurocognitive mechanisms of spontaneous language-based categorization of words. Importantly, our findings established a neural network of which dynamic activities underlie the processing of intra-language similarity and inter-language difference between words during language-based categorization of visual words. This network does not include the left middle and inferior frontal cortices which are fundamental for the processing of semantic, syntactic, and phonological information of words (Fedorenko et al., 2024a; Hodgson et al., 2021; Tan et al., 2005) but did not show neural RS effect related to language-based word categorization (see Fig. S18 for results). The word-categorization network is different from the language switching network that consists of the caudate (Crinion et al., 2006), lateral frontal cortices (Rodriguez-Fornells et al., 2005; Wang et al., 2007; Zhu et al., 2020), and anterior cingulate cortex (Abutalebi et al., 2007; Blanco-Elorrieta et al., 2018; Wang et al., 2007). The strength of the RS effects in the bilateral hub regions did not show dominance of one over the other hemisphere. This network includes the FULG in both hemispheres, unlike the visual word form area (VWFA) which predominates in the left hemisphere (Dehaene and Cohen, 2011; Zhan et al., 2023). While the PHC in the left hemisphere may contribute to verbal memory (Alessio et al., 2006), we showed that the PHC in both hemispheres were engaged in language-based categorization of visual words. The bilateral ATL and the LSTS are activated during comprehension of sentences requiring retrieval of information from autobiographical, emotional, and episodic memory (Ferstl et al., 2008). In our work, however, these regions were spontaneously involved during language-based word categorization in a perceptual task performed on a single word that did not demand any complicated memory processing. The insula and OFC can be activated by words of negative or positive valence (Alia-Klein et al., 2007; Gu and Han, 2007) or of object names with strong olfactory associations (Han et al., 2020).

The words used in our study, however, refer to names of tools or animals without obvious emotional contents. Our GCA results showed similar patterns of forward and backward connections between adjacent brain regions in the word-categorization network in both hemispheres, though the strength of connectivity between adjacent brain regions was slightly different between the two hemispheres.

The word-categorization network identified serves two key processes that support classification of words of two languages i.e., the computations of both intra-language similarity and inter-language difference between words. The neural processes of intra-language similarity and inter-language difference occurred spontaneously during language-based categorization of words because the one-back task employed in our study did not require explicit processing of linguistic properties of words or explicit classification of words of two different languages. However, the neural correlation distances corresponding to intra-language similarity and inter-language difference were highly correlated in the one-back and explicit classification tasks. Moreover, the neural index that combined intra-language similarity and inter-language difference (**δ**) predicted behavioral response efficiencies during an explicit word classification task. These results indicate that the two neural computations take place irrespective of task demands and provide a neural basis of behavioral performances during language-based word categorization.

Importantly, neural computations of intra-language similarity and inter-language difference occurred as early as 150 ms and completed within 300 ms after word onset, which were earlier than those involved in categorization of words in terms of semantic concepts (Giari et al., 2020) and different from the early word form processing that occurs dominantly in the left ventral occipito-temporal cortex around 170 ms after stimulus onset (Marinkovic et al., 2003; Nan et al., 2022; Pylkkänen and Marantz, 2003; Thesen et al., 2012). Other brain regions are engaged in a later time window (200–400 ms), including the left temporal cortex and ATL, and bilateral inferior prefrontal cortices and OFC (Marinkovic et al., 2003), which support semantic and phonological processing of words and sentences (Hodgson et al., 2021; Zhu et al., 2022). Our findings highlight rapid neural computations engaged in spontaneous language-based categorization of words which are independent of the processing of linguistic (e.g., semantic and phonological) information of words.

The word-categorization network serves spontaneous categorization of words of typologically diverse languages (i.e., Chinese, English, Korean, Italian) similarly in Chinese, English, and German speakers and for learned (e.g., Chinese and English) and unlearned languages (Korean and Italian). The human brain may evolve the network to enable spontaneous categorization of words of an alphabetic language and a non-alphabetic language as salient symbols of different social group identities. Indeed, some nodes of the word-categorization network have been found to respond to symbols of outgroup versus ingroup members (e.g., insula) (Merritt et al., 2021), underlie social categorization of faces and representations of social concepts (e.g., fusiform, ATLs) (Golby et al., 2001; Pobric et al., 2016; Zhou et al., 2020), and encode social information about others (e.g., OFC) (Park et al., 2020).

Chinese/Korean and English/Italian are languages that represent the two major cultural societies in the world (i.e., East Asian and Western Europe). Fast and spontaneous categorization of words of an alphabetic language and a non-alphabetic language as symbols of different cultural groups may provide a pivotal cognitive basis for classification of people for appropriate real-life social interactions.

Although the same neural network was engaged in the computations of intra-language similarity and inter-language difference during language-based word-categorization, the spatiotemporal characteristics of dynamic activities in this network underlying the two computations were not the same. The temporal procedures varied slightly across the two computations. The connectivity strength of the word-categorization network was greater during computations of intra-language similarity compared with inter-language difference. Moreover, the activations of the network started from different nodes during computations of intra-language similarity compared with inter-language difference. Therefore, although the essential function of the word-categorization network is to calculate a correlation distance between two words, this network may work in different fashions when computing the correlation distance between two words that belong to the same or different language categories.

How possible are the early neural RS effects within 200 ms after word onset observed in our study related to the processing of low-level perceptual features or high-level linguistic (e.g., orthography, semantics, phonology) properties of visual words? Our analyses of the ERPs to scrambled Chinese and English words in Experiment 2 did not show significant RS effect. Because only low-level visual features were preserved in the scrambled words, the ERP results provided no evidence that the early RS effects on the neural response to words can be attributed to habituation of perception of the low-level perceptual features. Furthermore, we found that the RS effects on the neural response to radicals and letters in Experiment 3 took place in a delayed time window and exhibited different scalp distributions (i.e., over the central region for radicals and occipital regions for letters) compared with the neural RS effects related to words. Thus the early RS effects on the neural response to words cannot be interpreted as habituation of perception of the middle-level units of Chinese and English words (i.e., radicals and letters) either. In addition, the early neural RS effects were similarly observed for both familiar (i.e., Chinese and English) and unfamiliar (i.e., Korean and Italian) languages and occurred earlier than the time window in which the processing of the linguistic properties of visual words takes place (Marinkovic et al., 2003; Hodgson et al., 2021; Zhu et al., 2022). Therefore, the early neural RS effects identified in our work were unlikely to be associated with the processing of the linguistic (e.g., orthography, semantics, phonology) properties of visual words since these properties of unfamiliar languages were unknown to the participants. Taken together, our findings of the early neural RS effects highlight an early word-level representation of alphabetic vs. non-alphabetic languages which distinguishes words from letters/radicals but is similar for familiar or unfamiliar languages. Our results, however, do not exclude the possibility that the processing of the linguistic properties of visual words may contribute to the long-latency RS effect around 300 ms after word onset. Further processing of the linguistic properties of visual words of familiar languages may follow the early language-based categorization of visual words, though this should be tested in future research.

Our findings raise other questions for future research. Our work tested only young adults. Given the finding of language-based social preferences in infants’ behaviors (Howard et al., 2015; Kinzler et al., 2012; Liberman et al., 2017a), it is interesting to compare the developmental trajectories of the neural underpinnings of the word-categorization network and the core linguistic language network. It is also important to clarify how these two networks interact with each other during language processing. The current work discovered the word-categorization network using only written words. It is unclear whether the key nodes of this network beyond fusiform/lingual gyri, which support word-form processing (Zhan et al., 2023), are also engaged in spontaneous language-based categorization of spoken words. To address this issue is important for understanding whether the spoken and writing systems of language function serve as markers of social group identities vis similar neural mechanisms. Recent research using single-neuronal recording has disentangled the process of semantic information (Jamali et al., 2024) and representations of internal and vocalized speech (Wandelt et al., 2024) at the cellular scale. Similar techniques may be employed to probe the mechanisms of language-based word categorization at the level of individual neurons in different nodes of the network. At last but not least, social group classification of others may occur based on both language signals and facial information (Rakić et al., 2011). How the word-categorization network interacts with that underlying social categorization of faces (Zhou et al., 2020) to influence real-life social behaviors deserves further investigation.

Finally, it should be noted that the current work was initiated by the previous behavioral findings which suggest that language can serve as a socially relevant category cue but focused on the neural mechanisms underlying rapid language-based categorization of visual words. Although the previous findings suggest that the language-based categorization of visual words provides a cognitive basis of social categorization of people, our work did not directly test whether and how the neural processes involved in the language-based categorization of visual words are linked to social evaluation or intergroup processes which are critical for social categorization of people. To clarify this issue should promote deep comprehension of the neural mechanisms underlying the social-categorization function of language but is beyond the scope of the current study. Future research should investigate the connection between language-based categorization of words and social categorization based on other social cues (e.g., faces), which is pivotal to understanding of social interactions in real-world situations.

In conclusion, our EEG and MEG results revealed robust RS effects in the early neural responses to visual words of the same language. The reliability of these RS effects was confirmed across words of different familiar and unfamiliar languages, in samples of speakers with different native languages, and through split-half reliability analyses (see Supplementary Materials, Fig. S19). These effects were supported by the bilateral neural networks whose activity reflected computations of correlation distances between word pairs, capturing both intra-language similarity and inter-language differences during the categorization of visual words in alphabetic and non-alphabetic languages. Together, these findings advance our understanding of spontaneous, language-based neural categorization of visual words as a key basis of the social-categorization function of language.

## Materials and Methods

### Participants

The present study recruited nine independent samples of native Chinese, English and German speakers in Experiments 1 to 9 (see Table S1 for information about participants). All participants were students recruited from universities in Beijing. All Chinese participants began to learn English in primary schools. The English speakers were from different regions (Experiment 4: 13 from North America, 10 from Southeast Asia, 7 from Europe, 1 from Australia, 1 from South Asia, 1 from the Middle East, and 1 from East Africa; Experiment 8: 23 from North America, 5 from Southeast Asia, 3 from Europe, 1 from Australia, 1 from South Asia and 1 from Hong Kong). All the German speakers in Experiment 5 were from Europe. All participants self-reported no neurological diagnoses and had normal or corrected-to-normal vision. This study was approved by the Research Ethics Committee at the School of Psychological and Cognitive Sciences, Peking University (Ethics approval number: #2022-02-07). Informed consent was obtained from all participants prior to the study. All participants were paid for their participation and were informed of their rights to quit at any time during the study. The sample size in Experiment 1 was determined in reference to a previous EEG/MEG study that investigated social categorization of faces using the same RS paradigm (Zhou et al., 2020). Sample sizes in the following studies were determined based on the results of Experiment 1.

### Stimuli

Language materials used in this study included 40 Chinese words, 40 English words and 40 German words (see Table S2 for all word stimuli). The two-character Chinese words included 20 animal names and 20 tool names. English, German, Korean, and Italian words were translated from the Chinese words to match semantic meanings. Word frequencies of these Chinese, English and German words were calculated using the SUBTLEX-CH (Cai and Brysbaert, 2010), SUBTLEX-US (Brysbaert and New, 2009), and SUBTLEX-DE (Brysbaert et al., 2011) databases.

Word frequencies (log10 frequency) were comparable for Chinese words (mean value = 0.97, from −0.62 to 2.74), English words (mean value = 1.32, from 0.09 to 2.68), and German words (mean value = 1.21, from −0.10 to 2.43). Each word subtended a visual angle of 11.53° × 4.28° at a viewing distance of 60 cm in EEG studies and of 8.67° × 3.88° at a view distance of 75 cm in MEG studies. Scrambled control stimuli were created by cutting each of the Chinese and English words into 32 squares (4 × 8) which were then shuffled and reorganized. This manipulation did not change the number of pixels and size of each stimulus. Twenty Chinese radicals and 20 English letters were selected. These stimuli were used to investigate potential contributions of shape features and middle-level units of words to language-based categorization of words.

### Procedure of the one-back task

The RS paradigm was adopted from previous studies of social categorization of faces (Zhang et al., 2023b; Zhou et al., 2020). Words of two languages (Chinese and English, English and German, or Korean and Italian), or scrambled words of two languages (Chinese and English), or word units of two languages (Chinese radicals and English letters) were used in different studies. This RS paradigm consisted of an alternating condition (Alt-Cond), in which words of two different languages (or scrambled words of two languages, or word units of two languages) were presented alternately, and a repetition condition (Rep-Cond), in which words of one language (or scrambled words of one language, or word units of one language) were presented repeatedly (Fig. 1a).

In each trial, a stimulus was displayed for 600 ms in the center of a grey background. Then, it was followed by a fixation cross which had a duration varying from 250 to 550 ms. Participants performed a one-back task (i.e., responding to a casual target stimulus that was presented in two consecutive trials) by pressing a button. In Experiments 1, 4 and 5, participants completed a Chinese-English session and a German-English session. The order of the two different sessions in each study was counter-balanced across participants. Each participant completed 3 runs in each session. Each run consisted of 8 blocks of 20 to 24 trials (including 20 non-target words and randomly 0 to 4 target words). A break of 8 s was given between two successive blocks in the way that a number count flashed with a 1-s step from 8 to 1 at the fixation position. In each run, words were presented in the Rep-Cond in 4 blocks of trials (two blocks for words of each language) and in the Alt-Cond in 4 blocks of trials. Words were displayed in a random order in each block and different blocks of trials were presented in a random order. This design gave 120 non-target trials of each language category in the Rep-Cond and Alt-Cond, respectively. In Experiments 7a and 8a, participants completed a Chinese-English session. The same design was employed in Experiment 2 using scrambled Chinese/English words and in Experiment 3 using Chinese radicals/English letters. The same design was used in Experiments 6 and 9 in which only Korean and Italian words were used.

### Procedure of the explicit word classification task

In Experiments 7b and 8b, participants were asked to perform a task to explicitly classify Chinese and English words during MEG recording. The stimuli (Chinese and English words) and procedure were the same as those in the Alt-Cond in Experiments 7a and 8a except that the inter-trial interval was 800 to 1400 ms. Participants were required to sort perceived words into Chinese or English by pressing two buttons as fast and accurately as possible. The response buttons corresponding to words of the two languages were counter-balanced across participants. Each participant completed 3 runs. Each run consisted of 40 Chinese words and 40 English words presented in a random order.

### EEG data acquisition and analyses

#### EEG data acquisition and preprocessing

We conducted EEG recordings using a 10-20 system cap with 64 Ag/AgCl ring electrodes (BrainAmp DC; Brain Products GmbH, Gilching, Germany). EEG signals were digitized at a sampling rate of 500 Hz with a band-pass filter of 0.01–100 Hz and referenced online against FCz. The electrode AFz was used as the ground.

Impedances of individual electrodes were kept below 5 kΩ. We performed EEG preprocessing and data analyses using the EEGLAB toolbox (Delorme and Makeig, 2004). The EEG signals were re-referenced to the average of the left and right mastoid electrodes (TP9, TP10) and filtered with a band-pass filter at 0.5–40 Hz during offline processing. After running independent component analysis, artifacts related to eye movement or blinks and muscle activities were automatically removed using the ICLabel (Pion-Tonachini et al., 2019) plug-in for EEGLAB with the criterion that probabilities of artifact components exceeded 0.9. Only non-target trials were included in EEG data analyses.

#### Univariate EEG data analyses

Event-related potentials (ERPs) in each condition were averaged separately with an epoch beginning 200 ms before stimulus onset and continuing for 1,000 ms. The baseline for measuring ERP amplitudes was the mean voltage of a 200-ms prestimulus interval. The latency of each ERP component was measured relative to stimulus onset. Trials with noise exceeding ±100 μV at any electrode were excluded from the average. This process left 118.94±3.00, 116.46±7.78, 118.59±3.57, 118.93±3.18, 117.73±5.16, and 119.65±0.83 trials on average in each condition for further EEG data analyses in Experiments 1 to 6, respectively. We used the FieldTrip toolbox (Oostenveld et al., 2011) to compare ERP amplitudes in the Rep-Cond and Alt-Cond. Cluster-based permutation *t*-tests were conducted to identify significant RS effects (i.e., decreased amplitudes in the Rep-Cond compared to Alt-Cond). A *t*-value indicating the difference in neural responses to words in the Rep-Cond and the Alt-Cond was calculated at each time point of each electrode. Adjacent points in time and space exceeding a predefined threshold (*P* < 0.05, two-tailed) were grouped into one or multiple clusters. The summed cluster *t*-values were compared against a permutation distribution to obtain the *P* values of each cluster. This distribution was generated by randomly reassigning condition markers for each participant (10,000 iterations) and the maximum summed cluster *t*-values were computed for each iteration (Maris and Oostenveld, 2007). We conducted two-tailed cluster-based permutation *t*-tests in the entire epoch (0–800 ms) at all electrodes (62 electrodes after excluding the reference channels (TP9 and TP10)) to identify significant clusters (*P* < 0.05) for all univariate EEG data analyses.

#### Multivariate EEG data analyses

Based on the results of univariate analyses that identified significant neural RS effects, we further conducted multivariate representation similarity analyses (Kriegeskorte et al., 2006) to examine the neural processes of intra-language similarity and inter-language difference engaged in spontaneous language-based categorization of words. We performed cluster-based permutation *t*-tests at the EEG electrodes which showed significant main effect of RS. A neural representation dissimilarity matrix (RDM) was constructed based on the correlation distance (one minus the Pearson correlation coefficients) between neural responses (ERP amplitudes at these electrodes) to two words from the same or different languages in both the Rep-Cond and Alt-Cond. All non-target trials were used in the computation of the RDM. This resulted in a 160 × 160 RDM, which included 40 words from each language in the Alt-Cond and Rep-Cond at each time point (Fig. 2c). Because the neural processes of intra-language similarity occurred more frequently in the Rep-Cond than Alt-Cond, the RDM corresponding to intra-language similarity would be attenuated in the Rep-Cond (vs. Alt-Cond) due to habituation (or the mean correlation distances would be increased in the Rep-Cond due to habituation). By contrast, the neural processes of inter-language difference occurred more frequently in the Alt-Cond than Rep-Cond, the RDM corresponding to inter-language difference would be attenuated in the Alt-Cond (vs. the Rep-Cond) due to habituation (or the mean correlation distances would be decreased in the Alt-Cond due to habituation). The differences in RDMs corresponding to both intra-language similarity and inter-language dissimilarity would be evident between the Rep-Cond and Alt-Cond if spontaneous language-based categorization of words engages both clustered representations of words of each language (i.e., enhanced processes of intra-language similarity in the Rep-Cond (vs. Alt-Cond)) and separated representations of words of two different languages (i.e., enhanced processes of inter-language dissimilarity or difference in the Alt-Cond (vs. Rep-Cond)). To test these predictions, we calculated the time courses of the mean values in the cells of the RDMs corresponding to intra-language dissimilarity and inter-language dissimilarity in both the Rep-Cond and Alt-Cond. After performing Fisher-Z transformation of the correlation distances, we then performed cluster-based permutation *t*-tests to examine significant difference between the time courses in 100 to 300 ms in the Rep-Cond and Alt-Cond (predefined threshold *P* < 0.025, 10,000 iterations, cluster-level *P* < 0.05, one-tailed).

After validating the time windows of significant multivariate RS effects on intra-language similarity and inter-language difference, we averaged these significant time periods to obtain three RDMs that represented the RS effects of intra-Chinese (Korean/Chinese radicals) similarity, intra-English (Italian/English letters) similarity, and Chinese vs. English (Korean vs. Italian/Chinese radicals vs. English letters) difference for each condition (the Rep-Cond or Alt-Cond). To further illustrate the RS effects of intra-language similarity and inter-language difference, with the MATLAB function *mdscale*, we performed non-metric multidimensional scaling analyses of the 160 × 160 RDM using Kruskal’s normalized stress formula-1 criterion. We extracted the first two dimensions and then transformed the RDM into three 2D space scatter plots for each condition. In this way, we could see how two kinds of language words are clustered within each language and separated from each other. The variations of clustered representations of words of the same language and separated representations of words of two languages were quantified by comparing the Euclidean distances in the word space between two words of the same language and between two words of different languages in the Alt-Cond and Rep-Cond, respectively.

### MEG data acquisition and analyses

#### MEG and MRI data acquisition and preprocessing

We used a whole-head MEG system with 102 magnetometers and 204 planar gradiometers (Elekta Neuromag TRIUX) to record neuromagnetic activity in a magnetically shielded room. The MEG signals were sampled at 1 kHz with an online band-pass filter of 0.1–330 Hz. We removed external interference from the raw MEG data using Maxfilter (Elekta-Neuromag) and co-registered head positions of each run with the first run using Maxmove (a subcomponent of Maxfilter). A high-resolution anatomical T1-weighted image was acquired for each participant (448× 512 mm matrix, 192 slices, 0.5 × 0.5 × 1.00 mm^3^ spatial resolution; TR = 2530 ms, TE=2.98 ms, inversion time (TI) = 1100 ms, FOV = 25.6 × 25.6 cm, FA = 7°, scanning order: interleaved). Padded clamps were used to minimize head motion and earplugs were used to attenuate scanner noise in the MRI acquisition. Three anatomical landmarks (nasion, left and right pre-auricular points), 4 HPI coils and at least 200 points on the scalp and face were digitized using the Probe Position Identification system (Polhemus) to co-register the MEG data with MRI coordinates. We conducted offline MEG preprocessing and analyses using the Brainstorm toolbox (Tadel et al., 2011). The MEG data were low-pass filtered at 40 Hz. Eye-blink artefacts were removed by signal-space projection. Then, the MEG data were epoched in terms of the stimulus trigger codes.

#### Sensor-space whole brain univariate analysis

Event-related fields (ERF) in each condition were averaged separately with an epoch beginning 200 ms before stimulus onset and continuing for 1,000 ms. The baseline for ERF measurement was the mean MEG activity of a 200-ms prestimulus interval and the latency was measured relative to stimulus onset. Trials exceeding 3500 fT at any MEG sensor were excluded from the average. This process left 118.27±3.49, 113.56±11.39, and 117.48±9.32 trials on average in each condition for further MEG data analyses in Experiments 7a, 8a, and 9, respectively. We examined the time courses of the RS effect on sensor-space MEG signals by pooling across Chinese and English words and Chinese and English speakers in Experiments 7a and 8a. Based on the EEG findings of neural RS effects, we conducted whole-brain cluster-based permutation *t*-tests to detect significant sensor-space RS effects (Alt-Cond > Rep-Cond, predefined threshold *P* < 0.01, 10,000 iterations, cluster-level *P* < 0.05, two-tailed) within 400 ms after stimulus onset.

#### Source-space whole brain and ROI univariate analyses

The high-resolution anatomical T1-weighted image of each participant was used for source reconstruction of the neural RS effect shown in sensor-space MEG signals. FreeSurfer (http://surfer.nmr.mgh.harvard.edu/) was used for segmentation of the T1 image. After co-registration of an individual’s brain anatomy and MEG sensors, the noise covariance matrix was computed from the 2-minute daily recordings of the empty room before the experiment. Brain activities were then estimated using a distributed model consisting of 15,002 current dipoles combining the time series of magnetometer and gradiometer signals using a linear inverse estimator (weighted minimum-norm current estimate, signal-to-noise ratio of 3, depth weighting of 0.5, unconstrained dipole orientations) separately for each condition and for each participant in a single-sphere head model. Individual source-space activities were then subject to baseline normalization by subtracting the mean and dividing by the standard deviation of source activations in the pre-stimulus intervals of 200 ms.

Normalized source activations were obtained by computing the norm of three dipole moments in each direction and were smoothed using an 8-mm FWHM Gaussian kernel. Individual normalized source-space data were projected to a standard brain model (ICBM152, 15,002 vertices) for the group-level analysis. Source space signals were down-sampled to 250 Hz before statistical analyses. The sources of neural RS effects were obtained by comparing the grand source signals across Chinese and English words in the Rep-Cond and Alt-Cond. Given the time window of significant RS effects on sensor-space signals, we performed whole-brain cluster-based permutation t-tests in 100–300 ms (vertex-level *P* < 0.001, 10,000 iterations, cluster-level *P* < 0.05, one-tailed) to detect significant attenuations of source-space MEG signals in the Rep-Cond than Alt-Cond. Based on the intersection between the RS effects in the source space and the Desikan-Killiany-Tourville atlas (Klein and Tourville, 2012), we created 11 regions of interest (ROI) to further assess the latency and intensity of RS effects in each brain region. Cluster-based permutation *t*-tests were performed in each brain region to testify significant RS effects (predefined *P* < 0.001, 10,000 iterations, cluster-level *P* < 0.05, one-tailed). We compared the peak intensities of the RS effects in the left and right hemisphere using 2 (left vs. right hemisphere) by 5 (OFC, insula, ATL, FULG, PHC region pairs) repeated measures analysis of variance (ANOVA) (LSTS was not included).

### Source-space multivariate analysis

We conducted multivariate analyses of source-space MEG signals to examine whether the neural network identified in the whole-brain univariate analyses underlies the neural processes of intra-language similarity and inter-language difference during language-based word categorization. Similar to the multivariate analyses of EEG signals, we computed a 160×160 RDM using MEG source space signals to Chinese and English words in the Rep-Cond and Alt-Cond. To increase the signal-to-noise ratio, we first subdivided the 11 ROIs into 313 subregions using the auto-subdivision function in Brainstorm (5 vertices in each subregion). We performed source reconstruction for each word in each condition without smoothing. The mean unsmoothed source signals of 1,000 Hz at each time point were extracted from these subregions to construct a 313-dimensional multivariate neural pattern for each word in each condition at each time point. Then, we computed a correlation distance (one minus the Pearson correlation coefficients) between patterns of MEG source signals of two words in the Rep-Cond and Alt-Cond to obtain the network 160×160 RDM at each time point. Correlation distances were calculated for two words of the same language and two words of two different languages as indices of intra-language similarity and inter-language difference, respectively. Cells in the RDM corresponding to intra-language similarity and inter-language difference were averaged to obtain the time course of the changes in correlation distances. Similarly, we conducted multidimensional scaling analyses of the global network RDMs to identify the first two components corresponding to each word. These two components were then used to construct a 2D word space in which words of the same language or of the two different languages in the Alt-Cond and Rep-Cond were plotted, respectively.

To further assess dynamic contributions of each brain region to the processing of intra-language similarity and inter-language difference during language-based word categorization, we computed local RDMs using the patterns of activity within each of the 11 ROIs. After performing Fisher-Z transformation, the mean correlation distance values specific to intra-/inter-language processing were calculated and assigned to all vertices in each ROI. This allowed us to get the whole-network correlation distance map in both the Rep-Cond and Alt-Cond for intra-language similarity and inter-language difference. Finally, we performed a whole-network cluster-based permutation *t*-test at 100–300 ms. A predefined threshold of *P* < 0.025, 10,000 iterations, and a cluster-level threshold of *P* < 0.05 (one-tailed) was used to examine the temporal and spatial characteristics of the neural RS effects related to the processing of intra-language similarity and inter-language difference.

#### Granger causality analysis

To further investigate information flow in the language categorization brain network, using the MVGC toolbox (Barnett and Seth, 2014), we conducted Granger causality analyses (GCA) (Geweke, 1984; Granger, 1969) of the time courses of the RS effects (i.e., the averaged correlation distance in the Rep-Cond minus that in the Alt-Cond at each time point during100–300 ms after the word onset) in the brain regions which demonstrated significant RS effects. The nodes of this network were the same as the MEG ROIs, including the bilateral OFC, insula, ATL, PHC, FULG, and the left STS.

Time series of intra-language similarity or inter-language difference were used to estimate the pairwise-conditional Granger causality among all brain regions. To satisfy the stationarity assumption and the requirement of zero-mean time series for GCA, we first preprocessed the time series of the RS effects, including linear detrending and rescaling (the subtraction of the temporal mean and division by the temporal standard deviation), to remove drifts and slow fluctuations of the time courses and to perform normalization. We conducted the preprocessing for all 780 samples (pairwise dissimilarity between all words of one language) for intra-language similarity and all of the 1600 samples (pairwise dissimilarity between all words of the two languages) for inter-language difference. We estimated the model order for each participant using the Bayesian information criterion and selected the model order of 5 (because down-sampling was not applied to the GCA analyses, a 5-ms lag was used for prediction of the neural activity in one brain region using the neural activity in another brain region) based on the mode of estimated model orders in all participants. A vector autoregression model was fitted using the time series data and was checked for stability and symmetric positive-definite residuals covariance matrix. All participants passed the model check. Then, we calculated time-domain pairwise-conditional causalities from the parameters of the vector autoregression model using the state-space method (Barnett and Seth, 2015). The vector autoregression model was transformed into an equivalent state-space model to estimate the Granger causality values (GC values). In this way, we obtained the Granger causality matrix for each participant and the averaged Granger causality matrix of all participants.

To test the group-level significance of the averaged Granger causality, we generated 1,000 surrogate averaged Granger causality matrices by stepwise permutation of the time series of the source variable by blocks (i.e., block permutation, randomly rearranging data while preserving local temporal dependencies). The original time series was divided into consecutive and non-overlapping blocks of fixed length. We used the model order, which is 5, as the block size. In each iteration, we permuted the time series of each variable while keeping the remaining variables unchanged to calculate the GC values from the permuted variable to the remaining variables. We conducted 1,000 iterations to obtain 1,000 surrogate averaged Granger causality matrices to make the null distribution of the Granger causality. Group-level significance of the original averaged Granger causality matrix was tested by comparing the original Granger causality with the null distribution. We employed the FDR method for correction for multiple comparisons because the GCA tested the connections among multiple brain regions. We reported and illustrated significant and reliable Granger causality connections in the language categorization brain network (*P* < 0.05, FDR correction, GC values > 0.001).

## Data availability

Data and codes for data analyses in this study are available at: http://datadryad.org/share/xB81CGC1LlwHF-D_G1A_fJJhvRSu5wnXs8jjG7lRjIo

## Acknowledgments

This work was supported by the National Natural Science Foundation of China (projects 32230043 and 32371092), Das Chinesisch-Deutsche Zentrum für Wissenschaftsförderung (M-0093), the High-performance Computing Platform of Peking University, and the National Center for Protein Sciences at Peking University. The authors thank Zhirui Zhao and Tengbin Huo for help with EEG and MEG data collection.

## Author contributions

SH and GZ conceived and designed the study, collected and analyzed the data, and wrote the manuscript. SH supervised all aspects of the research.

