## Supplementary Materials for "Neural categorization of visual words of alphabetic and non-alphabetic languages"

Figure S1-19

Table S1-5

### Supplementary Results

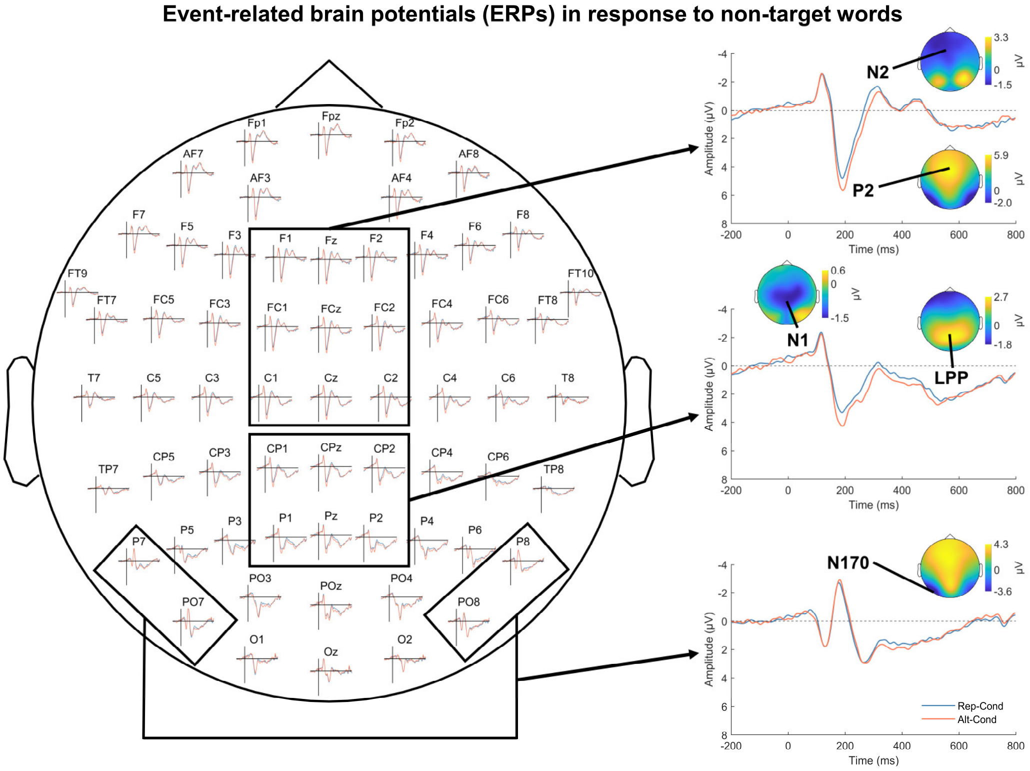


**Figure S1**. Illustrations of ERPs to non-target Chinese words at all electrodes in Chinese speakers in Experiment 1. The N170 showed the largest amplitude over the left occipito-temporal electrodes.

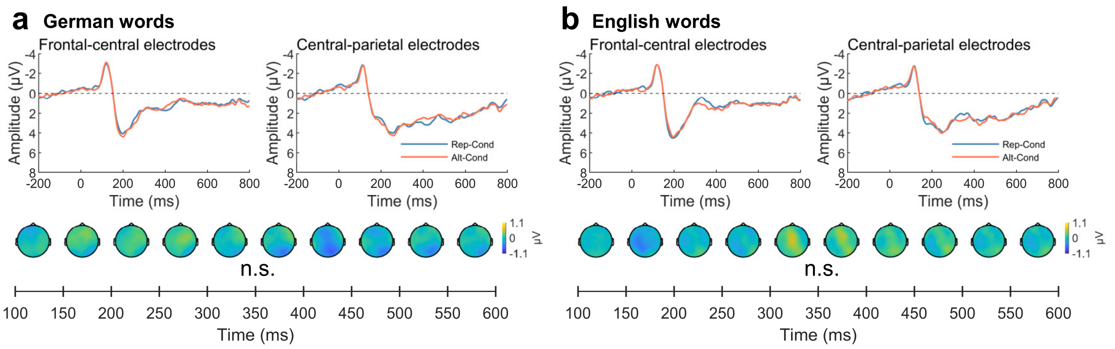


**Figure S2**. ERPs to German and English words in the Alt-Cond and Rep-Cond at the frontal-central and central parietal electrodes in Chinese speakers in Experiment 1. No significant difference was detected in the ERP amplitudes to English (or German) words between the Rep-Cond and Alt-Cond. n.s. = not significant.

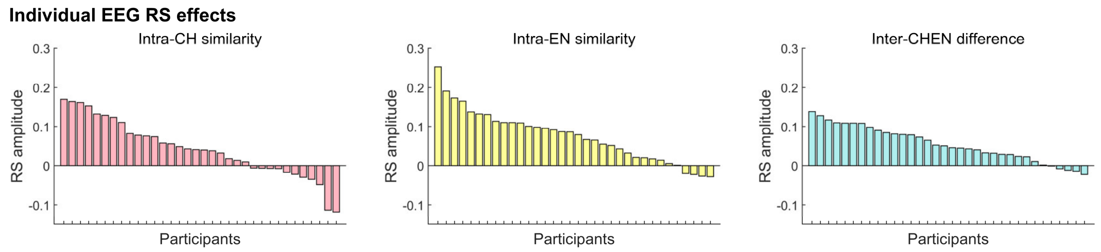


**Figure S3.** Illustration of the neural RS effects of each individual participant in the Chinese-English session of Experiment 1. The left and middle panels show the differences in the correlation distances of the RDMs corresponding to intra-language similarity between the Rep-Cond and Alt-Cond at 180–206 ms for Chinese words and at 168–226 ms for English words, respectively. The right panel shows the differences in the correlation distances of the RDMs corresponding to inter-language difference between the Alt-Cond and Rep-Cond at 162–218 ms. Intra-CH similarity = intra-Chinese word similarity; Intra-EN similarity = intra-English word similarity; Intra-CHEN difference = inter-Chinese/English word difference.

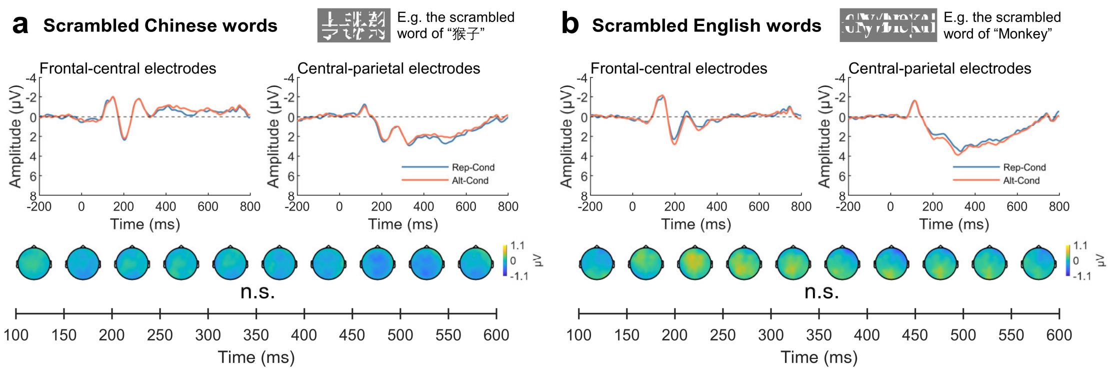


**Figure S4.** ERPs to scrambled Chinese and English words in the Alt-Cond and Rep-Cond at the frontal-central and central parietal electrodes in Chinese speakers in Experiment 2. Scramble words similarly elicited the N1/P2/N2/LPP components. However, there was no evidence for any significant RS effect on neural responses to scrambled Chinese or English words. n.s. = not significant.

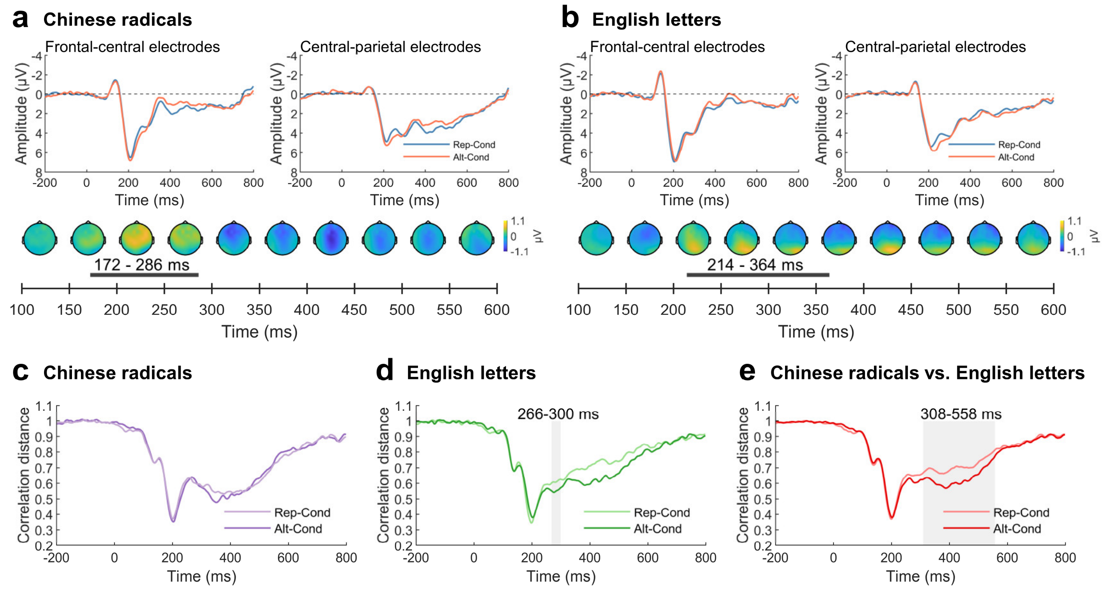


**Figure S5**. Results in Experiment 3. (a) and (b), ERPs to Chinese radicals and English letters in the Alt-Cond and Rep-Cond at the frontal-central and central-parietal electrodes in Chinese speakers in Experiment 3. (c), (d), (e), Results of the multivariate analyses.

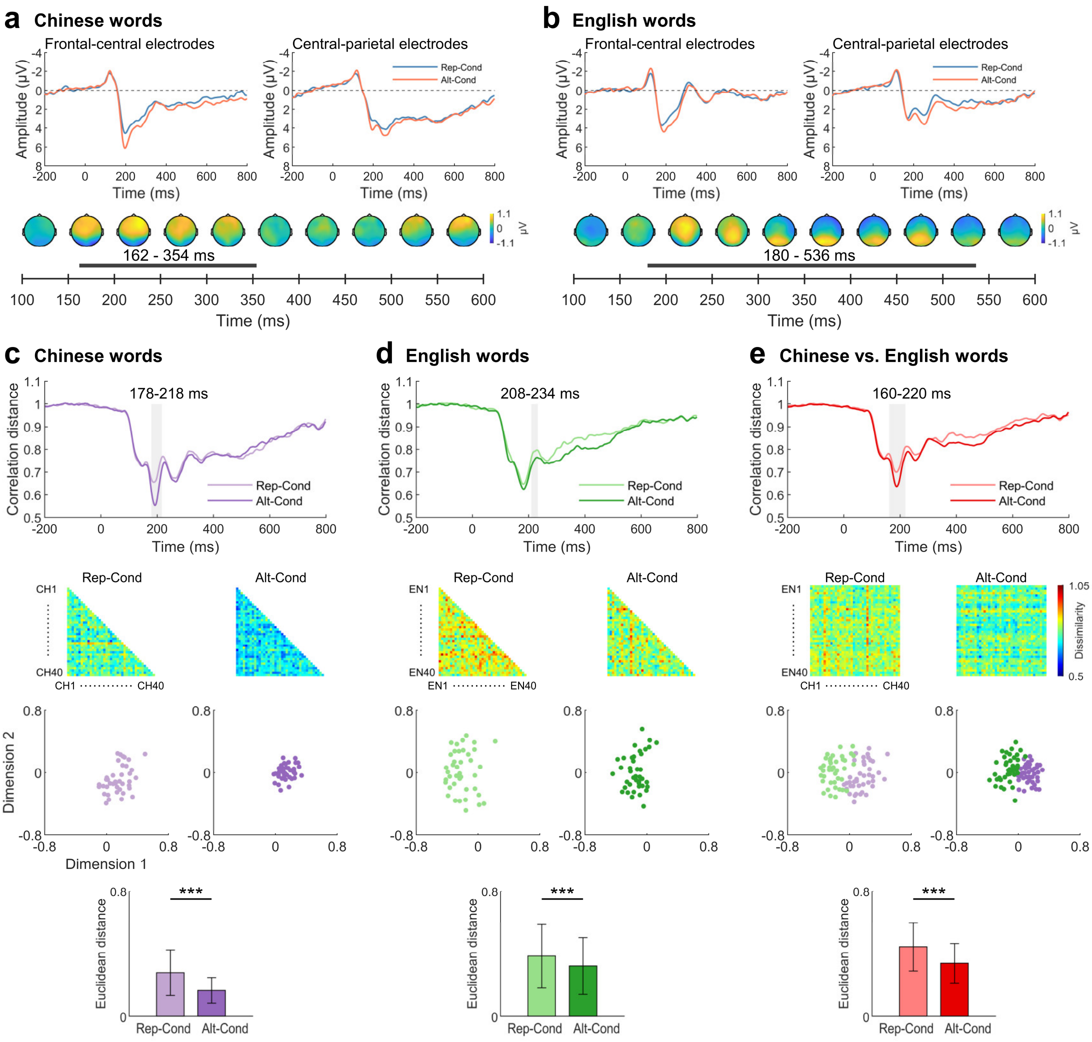


**Figure S6**. EEG results of English speakers in Experiment 4. (a) and (b) Results of univariate analyses. (c), (d), and (e) Results of multivariate and multidimensional scaling analyses. The top two panels show the time courses of significant differences in the correlation distances corresponding to intra-language similarity and inter-language difference between the Alt-Cond and Rep-Cond and the neural RDMs in the two conditions, respectively. The bottom two panels illustrate clustered representations of the words in the 2D word space based on the first two dimensions of multidimensional scaling analyses of the neural RDM corresponding to intra-language similarity and inter-language difference, respectively, and the mean Euclidean distances in the 2D word space between two words of the same language and between two words of different languages. *** $P < 0.001$.

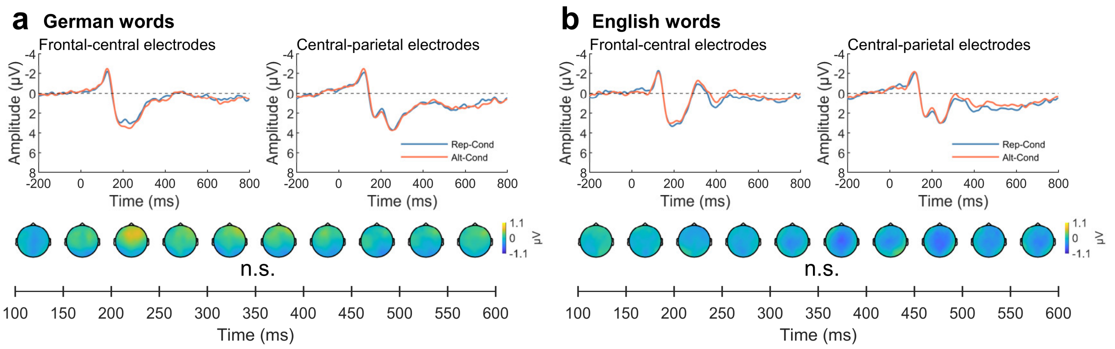


**Figure S7**. ERPs to German and English words in the Alt-Cond and Rep-Cond at the frontal-central and central-parietal electrodes in English speakers in Experiment 4. No significant RS effect was observed. n.s. = not significant.

In Experiment 4, univariate cluster-based permutation *t*-tests of the ERP amplitudes to non-target words in the Alt-Cond and Rep-Cond revealed significant RS effects in a cluster at 162–354 ms over the middle frontal/central/parietal regions for Chinese words. A significant cluster at 180–536 ms was identified for English non-target words which started from the middle frontal/central regions and then moved to the parietal and occipital regions (Fig. S6a and S6b). Similar analyses of ERPs to English and German words did not show any significant RS effect (Fig. S7). Further multivariate analyses of the neural responses to words showed a significantly increased correlation distance corresponding to intra-language similarity in the Rep-Cond (vs. Alt-Cond) at 178–218 ms for Chinese words and at 208–234 ms for English words (Fig. S6c-e). Moreover, the correlation distance corresponding to inter-language dissimilarity was significantly reduced in the Alt-Cond (vs. Rep-Cond) at 160–220 ms. Similarly, the multidimensional scaling analyses revealed more densely clustered representations of words of the same language in the Alt-Cond (vs. Rep-Cond) and more distantly separated representations of words of the two different languages in the Rep-Cond (vs. Alt-Cond) in the 2D word space. These results provide repeated evidence that the neural processes of intra-language similarity and inter-language difference were involved in spontaneous categorization of words between an alphabetic language (English) and a non-alphabetic (Chinese) language in English speakers.

Because the neural RS effects were observed for a native language and a non-native language in both Chinese and English speakers, it is critical to clarify whether similar neural RS effects manifesting language-based word categorization also occur between two non-native languages. We addressed this issue in Experiment 5 by recording EEG from a sample of native German speakers (N=34) who learned both Chinese and English. The stimuli and procedures were the same as those in Experiment 1. The results of both univariate and multivariate analyses replicated those in Experiments 1 and 4 (see Supplementary results, Fig. S8 and S9), indicating similar neural processes of intra-language similarity and inter-language difference involved in spontaneous categorization of words between an alphabetic language (English) and a non-alphabetic (Chinese) language but not between two alphabetic languages (English and German) in German speakers.

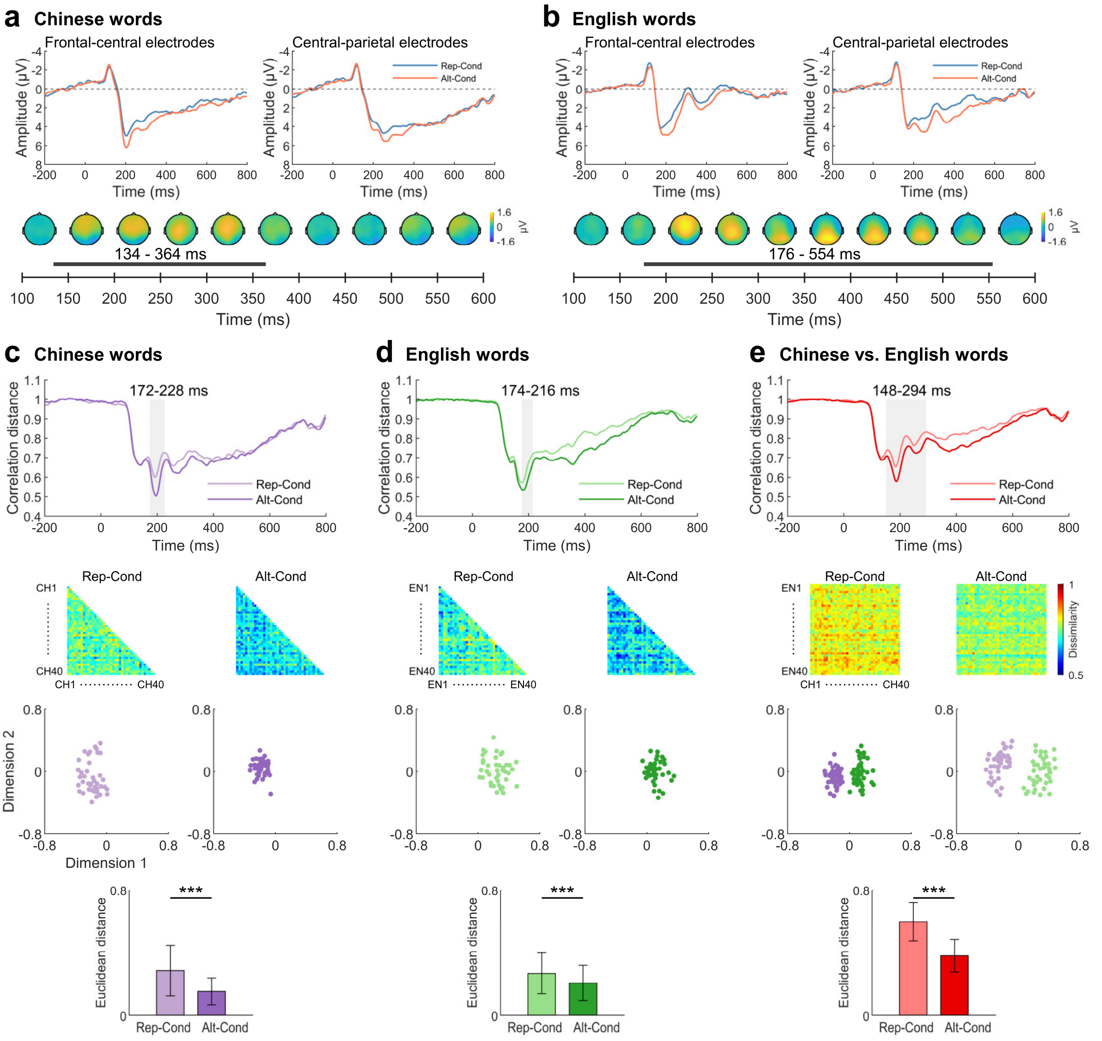


**Figure S8**. EEG results of German speakers in Experiment 5. (a) and (b) Results of univariate analyses. Top panels illustrate electrophysiological responses to Chinese and English words in the Alt-Cond and Rep-Cond, respectively. Bottom panels show voltage topographies of the scalp distributions of significant RS effects on neural responses to words of each language. (c), (d), and (e) Results of multidimensional scaling analyses. The top two panels show the time courses of significant differences in the correlation distances corresponding to intra-language similarity and inter-language difference between the Alt-Cond and Rep-Cond and the neural RDM in the two conditions, respectively. The bottom two panels illustrate clustered representations of words in the 2D word space built based on the first two dimensions of multidimensional scaling analyses of neural RDMs corresponding to intra-language similarity and inter-language difference, respectively, and the mean Euclidean distances in the 2D word space between two words of the same language and between two words of different languages.

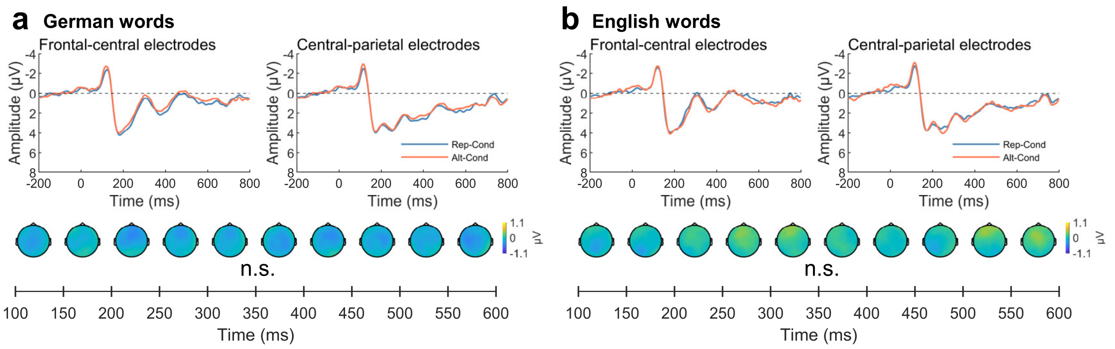


**Figure S9**. ERPs to German and English words in the Alt-Cond and Rep-Cond at the frontal-central and central-parietal electrodes in German speakers in Experiment 5. No significant RS effect was observed. n.s. = not significant.

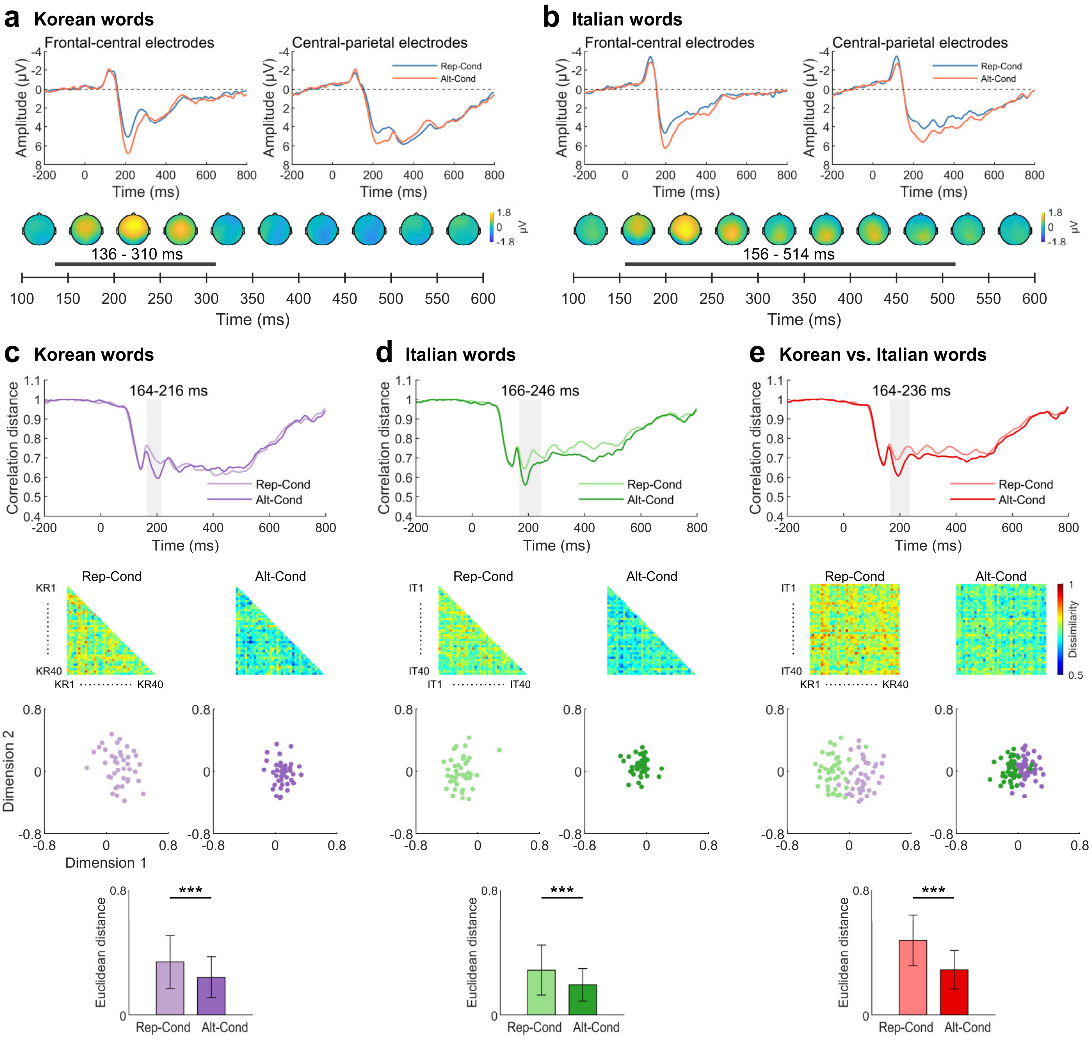


**Figure S10.** EEG results of Chinese speakers in Experiment 6. (a) and (b) Results of univariate analyses of electrophysiological responses to Korean and Italian words in the Alt-Cond and Rep-Cond, respectively. (c), (d), and (e) Results of multivariate and multidimensional scaling analyses. The top two panels show the time courses of significant differences in the correlation distances corresponding to intra-language similarity and inter-language difference between the Alt-Cond and the Rep-Cond and the neural RDMs in the two conditions, respectively. The bottom two panels illustrate clustered representations of the words in the 2D word space based on the first two dimensions of multidimensional scaling analyses of neural RDMs corresponding to intra-language similarity and inter-language difference, respectively, and the mean Euclidean distances in the 2D word space between two words of the same language and between two words of different languages. *** $P < 0.001$.

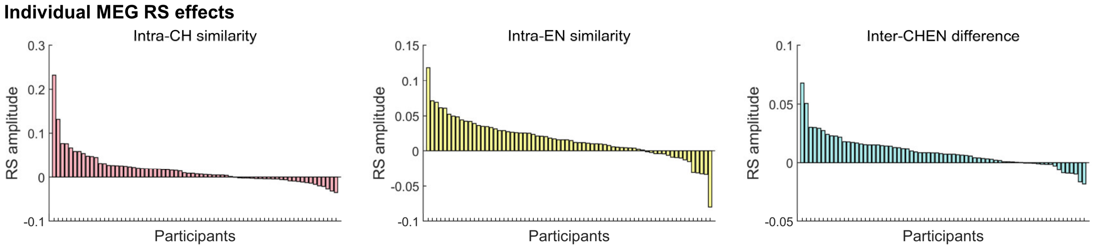


**Figure S11.** Illustration of the neural RS effects of each individual participant in Experiments 7a and 8a. The left and middle panels show the differences in the correlation distances of the RDMs corresponding to intra-language similarity between the Rep-Cond and Alt-Cond averaged at 159–200 ms for Chinese words and at 151–198 ms for English words, respectively. The right panel shows the differences in the correlation distances of the RDMs corresponding to inter-language difference between the Alt-Cond and Rep-Cond averaged at 150–232 ms and 236–294 ms. Intra-CH similarity = intra-Chinese word similarity; Intra-EN similarity = intra-English word similarity; Intra-CHEN difference = inter-Chinese/English word difference.

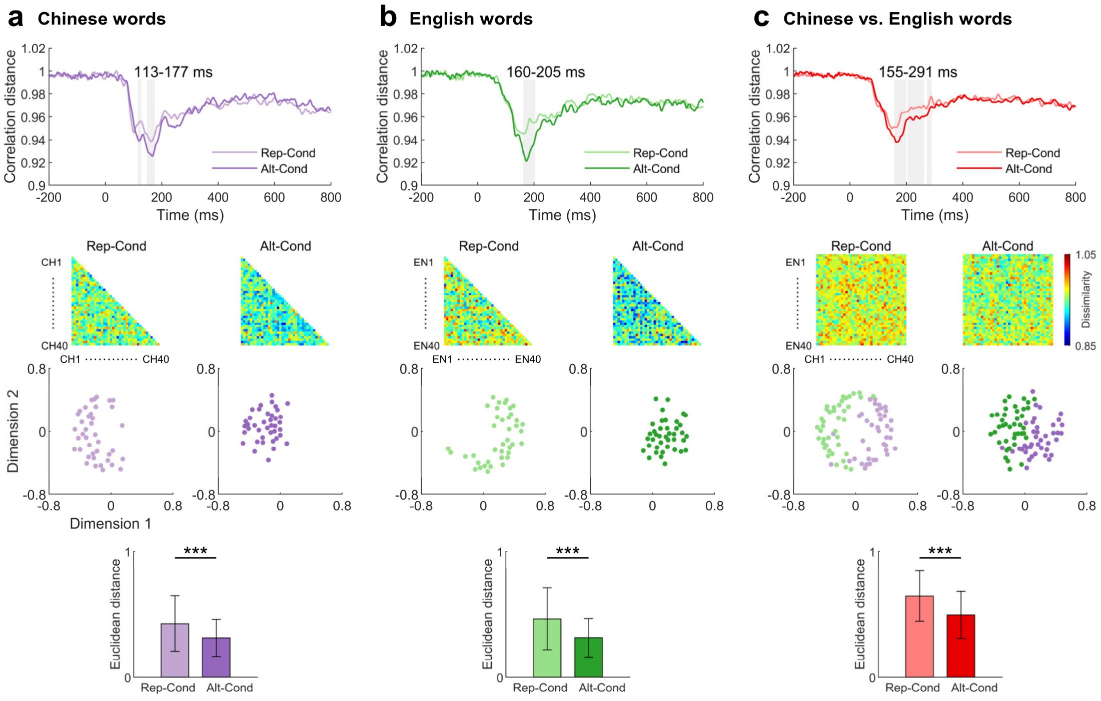


**Figure S12.** Results of the ROI analyses of MEG signals to Chinese or English words in Chinese speakers in Experiment 7a. The top two panels show the time courses of significant differences in the correlation distances corresponding to intra-language similarity and inter-language difference between the Alt-Cond and Rep-Cond, and the neural RDM in the two conditions, respectively. The bottom two panels illustrate clustered representations of words in the 2D word space built based on the first two dimensions of multidimensional scaling analyses of neural RDMs corresponding to intra-language similarity and inter-language difference, respectively, and the mean Euclidean distances in the 2D word space between two words of the same language and between two words of different languages. ***$P < 0.001$.

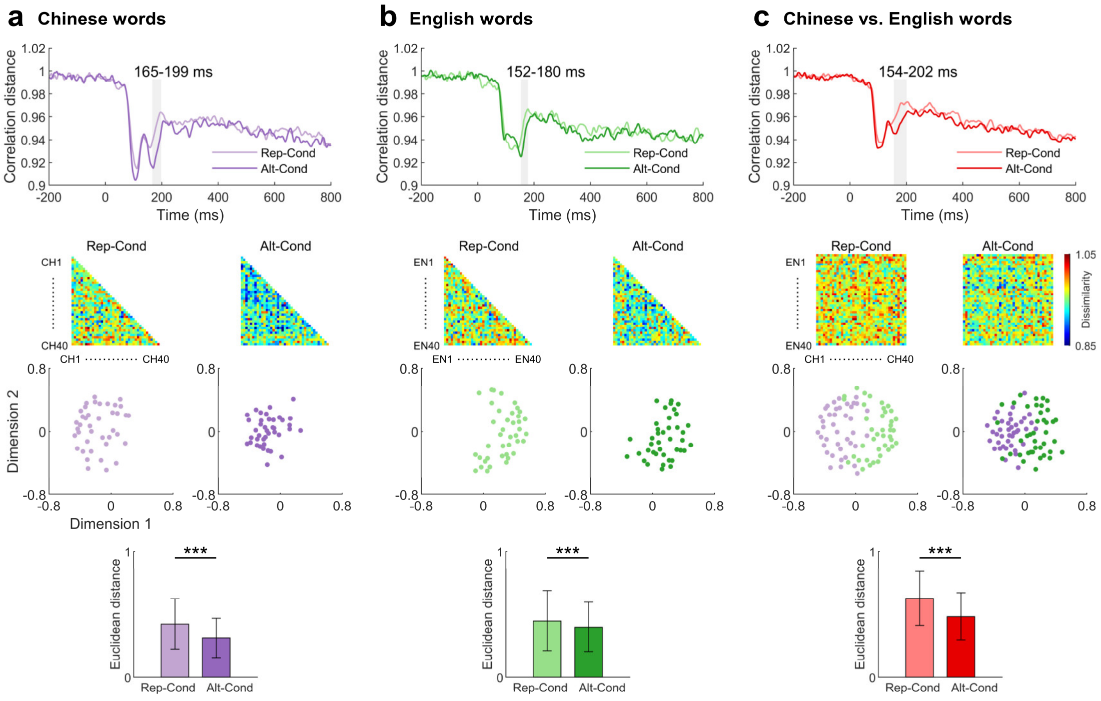


**Figure S13.** Results of the ROI analyses of MEG signals to Chinese or English words in English speakers in Experiment 8a. The top two panels show the time courses of significant differences in the correlation distances corresponding to intra-language similarity and inter-language difference between the Alt-Cond and Rep-Cond, and the neural RDM in the two conditions, respectively. The bottom two panels illustrate clustered representations of words in the 2D word space built based on the first two dimensions of multidimensional scaling analyses of neural RDMs corresponding to intra-language similarity and inter-language difference, respectively, and the mean Euclidean distances in the 2D word space between two words of the same language and between two words of different languages. ***$P < 0.001$.

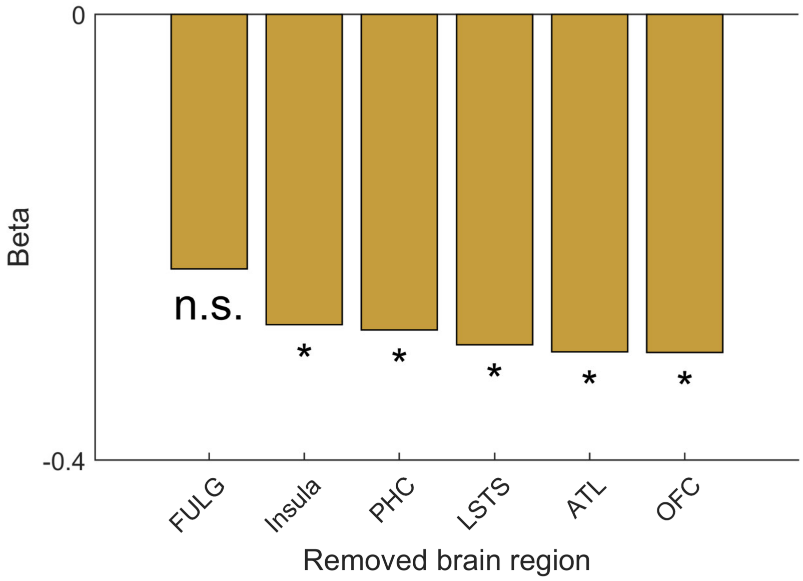


**Figure S14.** Results of the disruption analysis of the relationship between **δ** and IES. Shown are beta values of the regression analyses of the relationships between **δ** and behavioral categorization of words when the bilateral FULG, PHC, ATL, insula, OFC, and LSTS were removed from the network, respectively. The results showed that the association between **δ** and behavioral categorization of words was significant except when the bilateral FULGs were removed from the network. FULG=fusiform and lingual gyrus; PHC= parahippocampal cortex; STS=superior temporal sulcus; ATL=anterior temporal lobe; OFC=orbital frontal cortex. n.s.=no significance. $*P < 0.05$ (FDR correction).

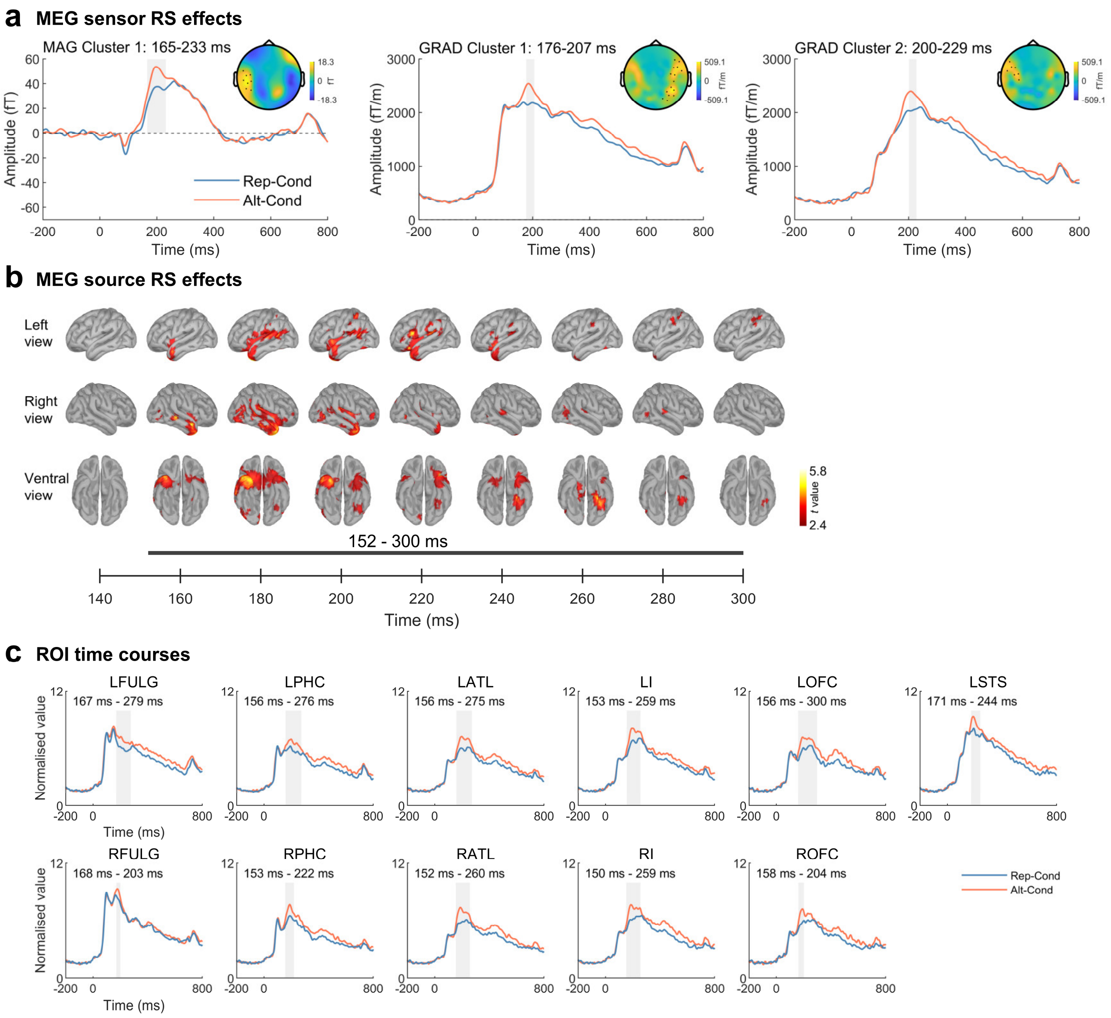


**Figure S15**. Results of whole-brain sensor-space and source-space MEG signals to Korean and Italian words in Chinese speakers in Experiment 9. Whole-brain RS effects of the source-space signals were identified using a lenient threshold (a predefined threshold of $P < 0.01$, 10,000 iterations, and a cluster-level threshold of $P < 0.05$, one-tailed). RS effects in the ROIs were identified using a lenient threshold (a predefined threshold of $P < 0.05$, 10,000 iterations, and a cluster-level threshold of $P < 0.05$, one-tailed).

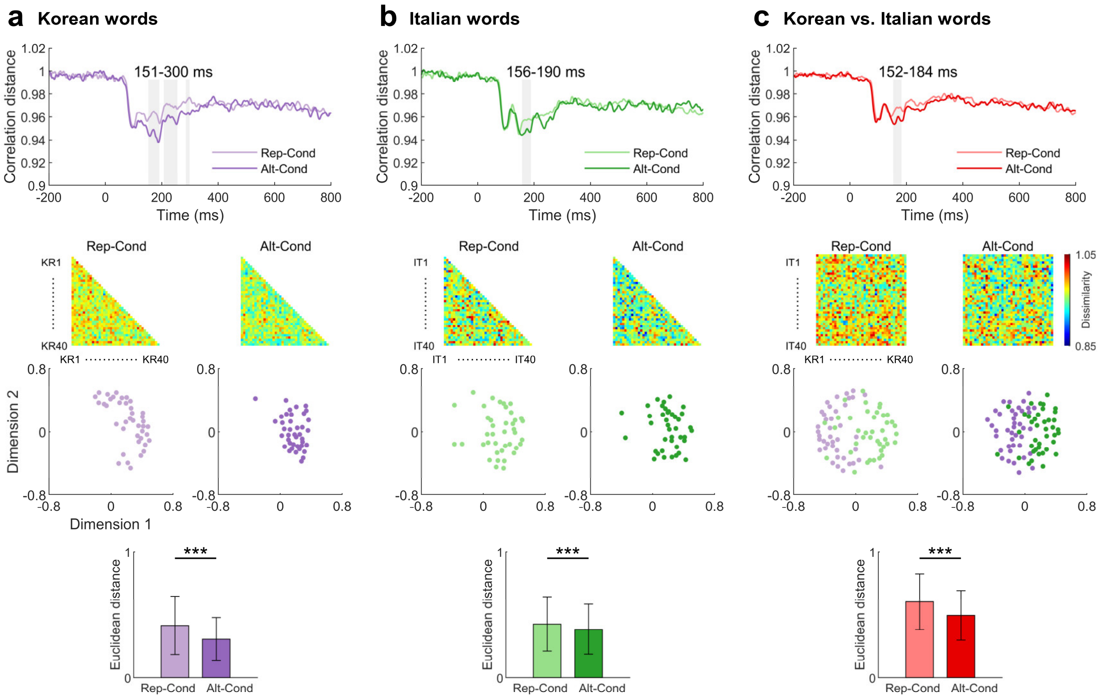

**Figure S16.** Results of the ROI analyses of MEG signals to Korean and Italian in Chinese speakers in Experiment 9. (a), (b), (c) Results of the ROI analyses of MEG signals to Korean or Italian words in native Chinese speakers in Experiment 9. The top two panels show the time courses of significant differences in the correlation distances corresponding to intra-language similarity and inter-language difference between the Alt-Cond and Rep-Cond, and the neural RDM in the two conditions, respectively. The bottom two panels illustrate clustered representations of words in the 2D word space built based on the first two dimensions of multidimensional scaling analyses of neural RDMs corresponding to intra-language similarity and inter-language difference, respectively, and the mean Euclidean distances in the 2D word space between two words of the same language and between two words of different languages. The RS effect of intra-language similarity of Italian words was identified using a lenient threshold (a predefined threshold of $P < 0.1$, 10,000 iterations, and a cluster-level threshold of $P < 0.05$, one-tailed) in the time window of 150–200 ms. *** P <0.001

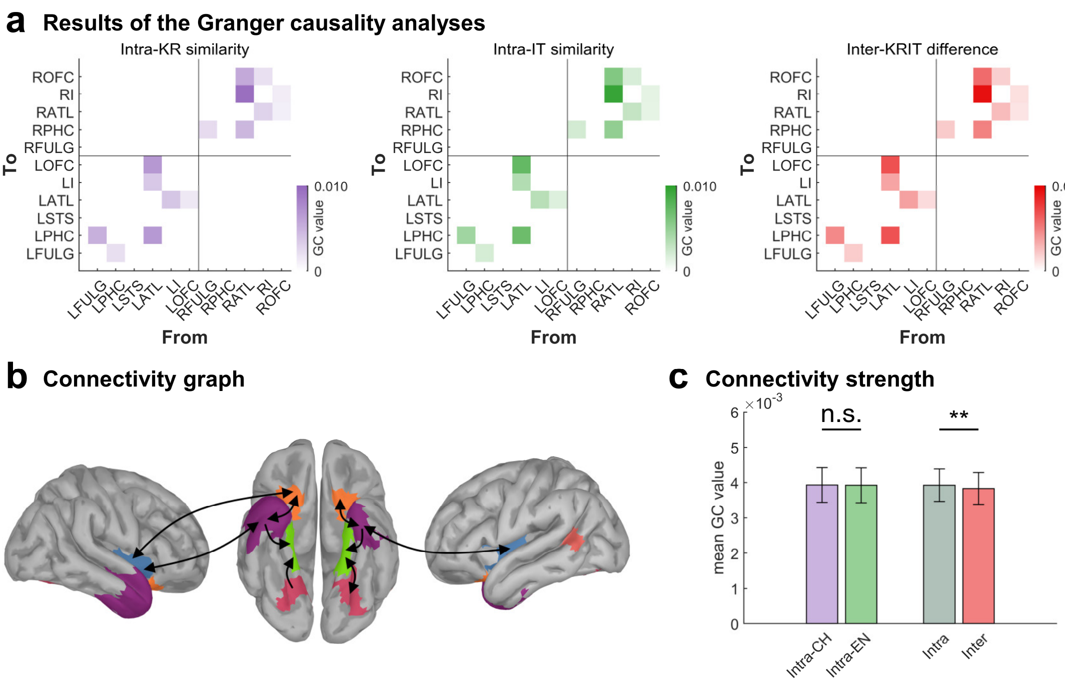


**Figure S17.** GCA results in Chinese speakers using Korean and Italian words in Experiment 9. Shown are (a) connectivity values above a threshold (GC values >0.001), (b) the connectivity patterns, and (c) the mean connectivity strength in the whole word-categorization network related to the processing of intra-language similarity and inter-language difference. ** $P$ <0.01

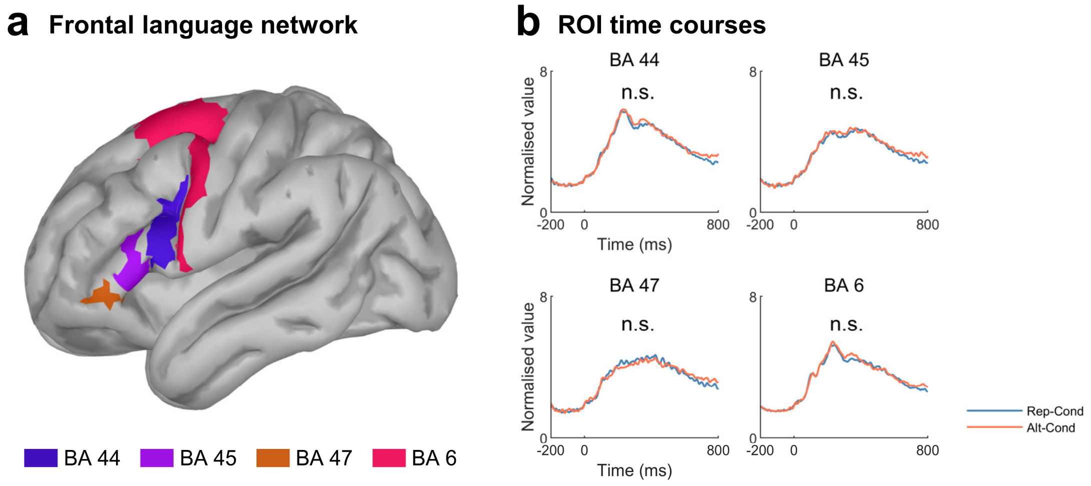


**Figure S18.** RS effects on neural responses to words in the frontal language network in Experiments 7a and 8a. (a) Illustration of the frontal language network that includes Brodmann areas 44, 45, 47, and 6. (b) Time courses of MEG source signals in each brain region of the frontal language network in the Alt-Cond and Rep-Cond. The Brodmann areas were defined based on the PALS atlas (Van Essen, 2005). BA 44 and 45 consist of the Broca's area as a key node of the domain-specific language network and domain-general multiple-demand network (Fedorenko and Blank, 2020). BA 47 belongs to the extended Broca's area and is involved in syntactic and semantic processing (Ardila et al., 2017; Friederici, 2011). BA 6 is associated with the integration of language and action during reading and naming tasks (Duffau et al., 2003; Pulvermüller, 2005). Source-space MEG signals from the Alt-Cond and Rep-Cond were plotted and subject to cluster-based permutation *t*-tests (predefined threshold of $P < 0.001$ and a cluster-level threshold of $P < 0.05$, one-tailed, 10,000 iterations). No significant RS effect was observed in the analyses that combined source-space MEG signals to Chinese and English words across Chinese and English speakers.

**Split-half reliability analysis of RDMs across participants**

We performed split-half reliability analyses of the neural RS effects in Experiment 1, 4, 5, 7 and 8 (Fig. S19). To quantify the stability of the representational structure over time, we estimated split-half reliability across participants in each experiment. At each time point, participants were randomly divided into two equal groups (17 vs. 17). For each group, the 160×160 RDMs were averaged across participants to obtain two group-level RDMs. To compute similarity between RDMs, only the upper triangular portion of each matrix (excluding the diagonal) was vectorized to avoid redundancy. The similarity between the two group-level RDMs was then quantified using Spearman rank correlation, providing a measure of the consistency of representational structure across independent subsets of participants. Because each split used only half of the available participants, correlation values were adjusted using the Spearman-Brown prophecy formula to estimate reliability at the full sample level. This procedure was repeated 200 times using different random splits of participants, yielding a distribution of reliability estimates for each time point. The mean across iterations was taken as the reliability estimate, and the standard deviation was computed to characterize variability due to random partitioning. This repeated split procedure provides a bootstrap-like estimate of reliability across participants. All analyses were conducted across a continuous time axis from -200 ms to 800 ms relative to stimulus onset.

To assess whether reliability values were significantly greater than zero, we performed a cluster-based permutation t test across time points. Statistical testing was restricted to post-stimulus time points (0-800 ms), as pre-stimulus activity was not expected to reflect stimulus-related representational structure. For each time point within this interval, the distribution of reliability estimates across iterations was compared against zero using a one-sided t test (testing whether the spearman rank correlation coefficient greater than zero). A predefined threshold of $P < 0.001$ was applied to identify candidate time points. Temporally contiguous significant time points were grouped into clusters, and a cluster-level statistic was computed for each cluster. A null distribution of cluster statistics was generated using 10,000 permutations, in which the data were randomly permuted across iterations to simulate the null hypothesis. Cluster-level *P*-values were obtained by comparing observed cluster statistics against this null distribution. Clusters with $P < 0.001$ were considered statistically significant.

For visualization, the mean split-half reliability across iterations was plotted as a function of time, with shaded regions indicating ±1 standard deviation across iterations. Significant temporal clusters identified by the permutation test were indicated as horizontal bars beneath the time course. The resulting reliability estimates reflect the consistency of representational geometry across independent subsets of participants. Given the limited number of trials per condition and the high dimensionality of the RDMs, reliability values are expected to be modest. Nevertheless, the reliability estimate is significantly greater than zero, indicating the presence of stable and reproducible representational structure over time.

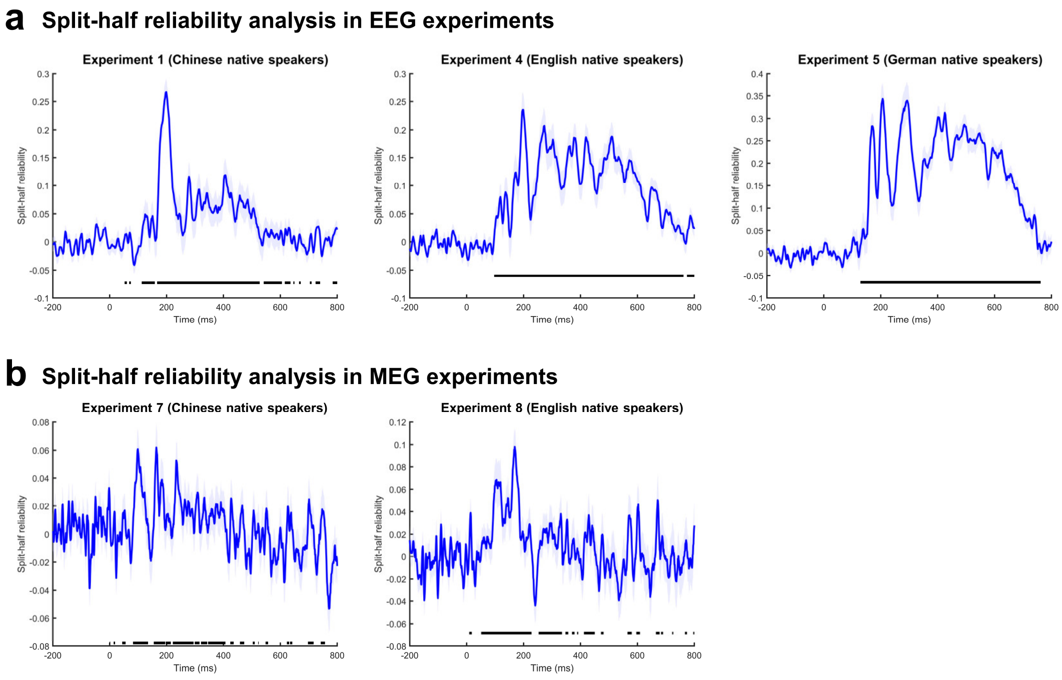


**Figure S19.** Time-resolved split-half reliability of the neural RDMs. (a) and (b) Mean split-half reliability (blue line) and ±1 SD (shaded) across 200 random splits in EEG and MEG experiments. Significance was assessed using a one-sided cluster-based permutation test against zero, restricted to 0-800 ms. Black bars indicate significant clusters greater than zero (predefined threshold of $P < 0.001$ and a cluster-level threshold of $P < 0.001$, 10,000 iterations).

**Table S1.** Demographic information about participants in Experiments 1−9

| Experiments | Sample size | Sex | Age (years) | Age range (years) | Chinese proficiency (years) | English proficiency (years) | German proficiency (years) |
|---|---|---|---|---|---|---|---|
| Experiment 1 | 34 | 17 males | 20.41±2.09 | 18~26 | native | 12.74±3.00 | / |
| Experiment 2 | 34 | 22 males | 21.21±2.69 | 17~24 | native | 12.65±3.13 | / |
| Experiment 3 | 34 | 13 males | 20.26±2.08 | 18~28 | native | 13.00±2.91 | / |
| Experiment 4 | 34 | 16 males | 22.15±2.12 | 19~28 | 9.12±6.67 | native | / |
| Experiment 5 | 34 | 20 males | 23.76±3.14 | 18~34 | 3.05±3.29 | 13.53±3.76 | native |
| Experiment 6 | 34 | 17 males | 19.85± 2.09 | 18~28 | native | 12.00±2.45 | / |
| Experiment 7a,7b | 34 | 17 males | 22.47±3.60 | 19~32 | native | 13.38±3.09 | / |
| Experiment 8a,8b | 34 | 17 males | 23.21±2.74 | 19~31 | 9.40±7.31 | native | / |
| Experiment 9 | 34 | 15 males | 20.35±2.06 | 17~25 | native | 12.56±2.68 | / |

*Notes.* The language proficiency was measured as the duration of formal language acquisition.

**Table S2.** Word stimuli used in Experiments 1, 4, 5, 6, 7, 8, and 9

| Chinese | English | German | Korean | Italian |
|---|---|---|---|---|
| 狮子 | Lion | Löwe | 사자 | Leone |
| 马匹 | Horse | Pferd | 말 | Cavallo |
| 山羊 | Goat | Ziege | 염소 | Capra |
| 鸭子 | Duck | Ente | 오리 | Anatra |
| 母牛 | Cow | Kuh | 소 | Mucca |
| 猴子 | Monkey | Affe | 원숭이 | Scimmia |
| 大象 | Elephant | Elefant | 코끼리 | Elefante |
| 猩猩 | Chimpanzee | Schimpanse | 침팬지 | Scimpanzé |
| 小鱼 | Fish | Fisch | 물고기 | Pesce |
| 狐狸 | Fox | Fuchs | 여우 | Volpe |
| 老鹰 | Eagle | Adler | 독수리 | Aquila |
| 小鹿 | Deer | Hirsch | 사슴 | Cervo |
| 野驴 | Donkey | Esel | 당나귀 | Asino |
| 袋鼠 | Kangaroo | Känguru | 캥거루 | Canguro |
| 黑熊 | Bear | Bär | 곰 | Orso |
| 青蛙 | Frog | Frosch | 개구리 | Rana |
| 小猫 | Cat | Katze | 고양이 | Gatto |
| 家犬 | Dog | Hund | 개 | Cane |
| 海豚 | Dolphin | Delfin | 돌고래 | Delfino |
| 公鸡 | Rooster | Hahn | 수탉 | Gallo |
| 电话 | Telephone | Telefon | 전화기 | Telefono |
| 钟表 | Clock | Uhr | 시계 | Orologio |
| 雨伞 | Umbrella | Regenschirm | 우산 | Ombrello |
| 汽车 | Car | Auto | 자동차 | Auto |
| 火车 | Train | Zug | 기차 | Treno |
| 地铁 | Subway | U-Bahn | 지하철 | Metropolitana |
| 飞机 | Airplane | Flugzeug | 비행기 | Aereo |
| 火箭 | Rocket | Rakete | 로켓 | Razzo |
| 胶水 | Glue | Leim | 접착제 | Colla |
| 桌子 | Table | Tisch | 테이블 | Tavolo |
| 椅子 | Chair | Stuhl | 의자 | Sedia |
| 按钮 | Button | Knopf | 단추 | Pulsante |
| 抽屉 | Drawer | Schublade | 서랍 | Cassetto |
| 冰箱 | Refrigerator | Kühlschrank | 냉장고 | Frigorifero |
| 钢笔 | Pen | Stift | 펜 | Penna |
| 尺子 | Ruler | Lineal | 자 | Righello |
| 书籍 | Book | Buch | 책 | Libro |
| 电视 | Television | Fernsehen | 텔레비전 | Televisione |
| 足球 | Football | Fußball | 축구공 | Calcio |
| 瓶子 | Bottle | Flasche | 병 | Bottiglia |

**Table S3.** Behavioral performances in the one-back task in Experiments 1–9

| | | | | |
|---|---|---|---|---|
| | **Experiment 1 (Chinese native speakers)** | | | |
| **Session** | **Chinese-English** | | | |
| Condition | Rep-CH | Rep-EN | Alt-CH | Alt-EN |
| Accuracy(%) | 96.08±0.07 | 97.06±0.05 | 98.23±0.04 | 96.07±0.07 |
| RT(ms) | 568.87±<br>62.93 | 577.87±<br>51.31 | 577.76±<br>70.90 | 579.90±<br>58.65 |
| IES | 597.86±<br>106.10 | 598.75±<br>78.67 | 589.87±<br>84.75 | 609.36±<br>101.43 |
| **Session** | **German-English** | | | |
| Condition | Rep-GE | Rep-EN | Alt-GE | Alt-EN |
| Accuracy(%) | 96.08±0.07 | 96.57±0.07 | 94.51±0.11 | 94.87±0.07 |
| RT(ms) | 567.51±<br>51.93 | 575.90±<br>65.83 | 570.94±<br>68.88 | 572.01±<br>58.33 |
| IES | 597.07±<br>97.49 | 603.69±<br>120.12 | 619.10±<br>155.78 | 607.88±<br>89.40 |
| | **Experiment 2 (Chinese native speakers)** | | | |
| **Session** | **Scrambled Chinese-English** | | | |
| Condition | Rep-SCH | Rep-SEN | Alt-SCH | Alt-SEN |
| Accuracy(%) | 87.99±0.12 | 85.29±0.11 | 85.15±0.15 | 87.35±0.09 |
| RT(ms) | 572.50±<br>45.55 | 591.29±<br>65.46 | 590.64±<br>69.20 | 575.52±<br>55.99 |
| IES | 664.56±<br>116.13 | 706.74±<br>136.33 | 737.82±<br>298.61 | 668.66±<br>115.21 |
| | **Experiment 3 (Chinese native speakers)** | | | |
| **Session** | **Chinese radicals-English letters** | | | |
| Condition | Rep-RD | Rep-LT | Alt-RD | Alt-LT |
| Accuracy(%) | 92.89±0.08 | 91.91±0.13 | 92.68±0.10 | 88.36±0.18 |
| RT(ms) | 586.79±<br>74.49 | 604.58±<br>79.74 | 595.52±<br>78.41 | 602.78±<br>95.71 |
| IES | 640.48±<br>125.79 | 681.05±<br>193.05 | 658.99±<br>170.99 | 786.51±<br>626.46 |
| | **Experiment 4 (English native speakers)** | | | |
| **Session** | **Chinese-English** | | | |
| Condition | Rep-CH | Rep-EN | Alt-CH | Alt-EN |
| Accuracy(%) | 94.12±0.09 | 94.36±0.08 | 93.79±0.08 | 95.22±0.07 |
| RT(ms) | 568.62±<br>68.39 | 553.12±<br>70.98 | 572.36±<br>69.51 | 563.14±<br>76.27 |
| IES | 613.65±<br>123.58 | 594.68±<br>121.63 | 618.07±<br>116.99 | 595.45±<br>101.52 |
| **Session** | **German-English** | | | |
| Condition | Rep-GE | Rep-EN | Alt-GE | Alt-EN |
| Accuracy(%) | 93.63±0.07 | 94.36±0.09 | 93.87±0.09 | 95.29±0.07 |
| RT(ms) | 565.77± | 562.26± | 571.89± | 560.00± |

| | | | | |
|---|---|---|---|---|
| | 71.28 | 74.15 | 67.14 | 78.80 |
| IES | 610.38± 104.48 | 603.51± 113.77 | 616.59± 103.86 | 592.48± 106.46 |
| | **Experiment 5 (German native speakers)** | | | |
| **Session** | **Chinese-English** | | | |
| Condition | Rep-CH | Rep-EN | Alt-CH | Alt-EN |
| Accuracy(%) | 89.71±0.10 | 91.67±0.12 | 92.95±0.09 | 93.04±0.10 |
| RT(ms) | 583.35± 60.74 | 575.23± 74.58 | 582.86± 68.93 | 578.84± 60.55 |
| IES | 660.29± 121.25 | 642.70± 145.40 | 633.36± 105.32 | 633.51± 125.41 |
| **Session** | **German-English** | | | |
| Condition | Rep-GE | Rep-EN | Alt-GE | Alt-EN |
| Accuracy(%) | 93.38±0.10 | 93.38±0.11 | 92.61±0.09 | 91.99±0.12 |
| RT(ms) | 579.84± 84.87 | 574.56± 71.39 | 573.63± 85.37 | 579.65± 78.41 |
| IES | 637.01± 175.36 | 632.08± 168.75 | 628.80± 138.81 | 655.02± 204.02 |
| | **Experiment 6 (Chinese native speakers)** | | | |
| **Session** | **Korean-Italian** | | | |
| Condition | Rep-KR | Rep-IT | Alt-KR | Alt-IT |
| Accuracy(%) | 91.42±0.09 | 95.83±0.06 | 91.01±0.10 | 94.92±0.08 |
| RT(ms) | 567.54± 63.37 | 571.46± 47.05 | 557.62± 60.88 | 581.88± 51.57 |
| IES | 627.45± 97.54 | 597.98± 55.39 | 619.77± 94.45 | 618.14± 86.30 |
| | **Experiment 7a (Chinese native speakers)** | | | |
| **Session** | **Chinese-English** | | | |
| Condition | Rep-CH | Rep-EN | Alt-CH | Alt-EN |
| Accuracy(%) | 94.61±0.08 | 92.89±0.10 | 95.03±0.07 | 92.47±0.07 |
| RT(ms) | 583.01± 69.17 | 585.27± 68.04 | 590.05± 70.82 | 591.92± 68.54 |
| IES | 621.91± 97.81 | 641.58± 133.50 | 625.19± 94.31 | 646.45± 112.91 |
| | **Experiment 8a (English native speakers)** | | | |
| **Session** | **Chinese-English** | | | |
| Condition | Rep-CH | Rep-EN | Alt-CH | Alt-EN |
| Accuracy(%) | 92.16±0.09 | 97.06±0.05 | 95.64±0.07 | 95.50±0.06 |
| RT(ms) | 580.11± 69.21 | 566.32± 63.93 | 574.91± 56.82 | 551.50± 64.55 |
| IES | 640.99± 131.84 | 585.73± 79.17 | 605.90± 88.05 | 581.03± 88.65 |
| | **Experiment 9 (Chinese native speakers)** | | | |
| **Session** | **Korean-Italian** | | | |

| Condition | Rep-KR | Rep-IT | Alt-KR | Alt-IT |
|---|---|---|---|---|
| Accuracy(%) | 92.65±0.07 | 94.12±0.08 | 89.57±0.11 | 94.78±0.09 |
| RT(ms) | 564.02±<br>58.15 | 570.92±<br>68.39 | 554.83±<br>67.08 | 563.77±<br>52.02 |
| IES | 614.21±<br>92.16 | 613.14±<br>108.48 | 632.32±<br>138.80 | 602.62±<br>98.28 |

*Notes.* CH, Chinese; EN, English; GE, German; SCH, Scrambled Chinese; SEN, Scrambled English; RD, Radicals; LT, Letters; KR, Korean; IT, Italian; IES, Inverse efficiency score.

**Table S4.** Statistical details of cluster-based permutation t tests in Experiments 1–9

| | | | |
|---|---|---|---|
| | | **Experiment 1** | |
| **Session** | | **Chinese-English** | |
| **Analysis** | | **Univariate analysis** | |
| Condition | Alt-CH vs. Rep-CH | Alt-EN vs. Rep-EN | Alt-Cond vs. Rep-Cond |
| Predefined *P* value | < 0.05 | < 0.05 | < 0.05 |
| Cluster-level *P* value | < 0.001 | < 0.001 | < 0.001 |
| **Analysis** | | **Multivariate analysis** | |
| Condition | Alt-CH vs. Rep-CH | Alt-EN vs. Rep-EN | Alt-CHEN vs. Rep-CHEN |
| Predefined *P* value | < 0.025 | < 0.025 | < 0.025 |
| Cluster-level *P* value | 0.025 | < 0.001 | < 0.001 |
| **Session** | | **German-English** | |
| **Analysis** | | **Univariate analysis** | |
| Condition | Alt-GE vs. Rep-GE | Alt-EN vs. Rep-EN | |
| Predefined *P* value | < 0.05 | < 0.05 | |
| Cluster-level *P* value | 0.672 | 0.061 | |
| | | **Experiment 2** | |
| **Session** | | **Scrambled Chinese-English** | |
| **Analysis** | | **Univariate analysis** | |
| Condition | Alt-SCH vs. Rep-SCH | Alt-SEN vs. Rep-SEN | |
| Predefined *P* value | < 0.05 | < 0.05 | |
| Cluster-level *P* value | 1 | 0.144 | |
| | | **Experiment 3** | |
| **Session** | | **Chinese radicals-English letters** | |
| **Analysis** | | **Univariate analysis** | |
| Condition | Alt-RD vs. Rep-RD | Alt-LT vs. Rep-LT | Alt-Cond vs. Rep-Cond |
| Predefined *P* value | < 0.05 | < 0.05 | < 0.05 |
| Cluster-level *P* value | 0.021 | 0.026 | 0.009 |
| **Analysis** | | **Multivariate analysis** | |

| Condition | Alt-RD vs. Rep-RD | Alt-LT vs. Rep-LT | Alt-RDLT vs. Rep-RDLT |
| --- | --- | --- | --- |
| Predefined *P* value | < 0.025 | < 0.025 | < 0.025 |
| Cluster-level *P* value | 1 | 0.009 | < 0.001 |
| | | **Experiment 4** | |
| **Session** | | **Chinese-English** | |
| **Analysis** | | **Univariate analysis** | |
| Condition | Alt-CH vs. Rep-CH | Alt-EN vs. Rep-EN | Alt-Cond vs. Rep-Cond |
| Predefined *P* value | < 0.05 | < 0.05 | < 0.05 |
| Cluster-level *P* value | < 0.001 | < 0.001 | < 0.001 |
| **Analysis** | | **Multivariate analysis** | |
| Condition | Alt-CH vs. Rep-CH | Alt-EN vs. Rep-EN | Alt-CHEN vs. Rep-CHEN |
| Predefined *P* value | < 0.025 | < 0.025 | < 0.025 |
| Cluster-level *P* value | 0.005 | 0.021 | < 0.001 |
| **Session** | | **German-English** | |
| **Analysis** | | **Univariate analysis** | |
| Condition | Alt-GE vs. Rep-GE | Alt-EN vs. Rep-EN | |
| Predefined *P* value | < 0.05 | < 0.05 | |
| Cluster-level *P* value | 0.131 | 0.332 | |
| | | **Experiment 5** | |
| **Session** | | **Chinese-English** | |
| **Analysis** | | **Univariate analysis** | |
| Condition | Alt-CH vs. Rep-CH | Alt-EN vs. Rep-EN | Alt-Cond vs. Rep-Cond |
| Predefined *P* value | < 0.05 | < 0.05 | < 0.05 |
| Cluster-level *P* value | < 0.001 | < 0.001 | < 0.001 |
| **Analysis** | | **Multivariate analysis** | |
| Condition | Alt-CH vs. Rep-CH | Alt-EN vs. Rep-EN | Alt-CHEN vs. Rep-CHEN |
| Predefined *P* value | < 0.025 | < 0.025 | < 0.025 |

| | | | |
|---|---|---|---|
| Cluster-level *P* value | < 0.001 | 0.004 | < 0.001 |
| **Session** | | **German-English** | |
| **Analysis** | | **Univariate analysis** | |
| Condition | Alt-GE vs. Rep-GE | Alt-EN vs. Rep-EN | |
| Predefined *P* value | < 0.05 | < 0.05 | |
| Cluster-level *P* value | 0.634 | 0.310 | |
| | | **Experiment 6** | |
| **Session** | | **Korean-Italian** | |
| **Analysis** | | **Univariate analysis** | |
| Condition | Alt-KR vs. Rep-KR | Alt-IT vs. Rep-IT | Alt-Cond vs. Rep-Cond |
| Predefined *P* value | < 0.05 | < 0.05 | < 0.05 |
| Cluster-level *P* value | < 0.001 | < 0.001 | < 0.001 |
| **Analysis** | | **Multivariate analysis** | |
| Condition | Alt-KR vs. Rep-KR | Alt-IT vs. Rep-IT | Alt-KRIT vs. Rep-KRIT |
| Predefined *P* value | < 0.025 | < 0.025 | < 0.025 |
| Cluster-level *P* value | 0.002 | < 0.001 | < 0.001 |
| | | **Experiment 7a and 8a** | |
| **Session** | | **Chinese-English** | |
| **Analysis** | | **Univariate analysis (sensor space)** | |
| Condition | | Alt-Cond vs. Rep-Cond | |
| Predefined *P* value | | < 0.01 | |
| Cluster-level *P* value | MAG Cluster 1 *P* < 0.001 | MAG Cluster 2 *P* < 0.001 | GRAD Cluster 1 *P* < 0.001 |
| **Analysis** | | **Univariate analysis (source space)** | |
| Condition | | Alt-Cond vs. Rep-Cond | |
| Predefined *P* value | | < 0.001 | |
| Cluster-level *P* value | Cluster 1 *P* < 0.001 | Cluster 2 *P* < 0.001 | Cluster 3 *P* = 0.020 |
| **Analysis** | | **ROI analysis** | |
| Condition | | Alt-Cond vs. Rep-Cond | |
| Predefined *P* value | | < 0.001 | |
| Cluster-level *P* value | | All P values of clusters in the ROIs < 0.003 | |

| Analysis | Multivariate analysis (7a and 8a) | | |
|---|---|---|---|
| Condition | Alt-CH vs. Rep-CH | Alt-EN vs. Rep-EN | Alt-CHEN vs. Rep-CHEN |
| Predefined *P* value | < 0.025 | < 0.025 | < 0.025 |
| Cluster-level *P* value | 0.003 | 0.001 | < 0.001 |
| **Analysis** | **Multivariate analysis (7a)** | | |
| Condition | Alt-CH vs. Rep-CH | Alt-EN vs. Rep-EN | Alt-CHEN vs. Rep-CHEN |
| Predefined *P* value | < 0.025 | < 0.025 | < 0.025 |
| Cluster-level *P* value | Cluster 1 $P = 0.011$; Cluster 2 $P = 0.042$; | 0.001 | Cluster 1 and 2 $P < 0.001$; Cluster 3 $P = 0.034$ |
| **Analysis** | **Multivariate analysis (8a)** | | |
| Condition | Alt-CH vs. Rep-CH | Alt-EN vs. Rep-EN | Alt-CHEN vs. Rep-CHEN |
| Predefined *P* value | < 0.025 | < 0.025 | < 0.025 |
| Cluster-level *P* value | 0.011 | 0.014 | 0.002 |
| | **Experiment 9** | | |
| **Session** | **Korean-Italian** | | |
| **Analysis** | **Univariate analysis (sensor space)** | | |
| Condition | Alt-Cond vs. Rep-Cond | | |
| Predefined *P* value | < 0.01 | | |
| Cluster-level *P* value | MAG Cluster 1 $P = 0.018$ | GRAD Cluster 1 $P < 0.001$ | GRAD Cluster 2 $P = 0.018$ |
| **Analysis** | **Univariate analysis (source space)** | | |
| Condition | Alt-Cond vs. Rep-Cond | | |
| Predefined *P* value | < 0.01 | | |
| Cluster-level *P* value | Cluster 1 and Cluster 2 $P < 0.001$; Cluster 3 $P = 0.030$; Cluster 4 $P = 0.037$ | | |
| **Analysis** | **ROI analysis** | | |
| Condition | Alt-Cond vs. Rep-Cond | | |
| Predefined *P* value | < 0.05 | | |
| Cluster-level *P* value | All P values of clusters in the ROIs < 0.036 | | |
| **Analysis** | **Multivariate analysis** | | |
| Condition | Alt-KR vs. Rep-KR | Alt-IT vs. Rep-IT | Alt-KRIT vs. Rep-KRIT |
| Predefined *P* | < 0.025 | < 0.025 | < 0.025 |

| | | | |
|---|---|---|---|
| value | | | |
| Cluster-level *P* value | Cluster 1 and 2 $P < 0.001$; Cluster 3 $P = 0.041$ | 0.021 | 0.006 |

*Notes.* CH, Chinese; EN, English; GE, German; SCH, Scrambled Chinese; SEN, Scrambled English; RD, Radicals; LT, Letters; KR, Korean; IT, Italian.

**Table S5.** Statistical details of t tests in multivariate analysis

| | | | |
|---|---|---|---|
| | | **Experiment 1** | |
| Condition | Alt-CH vs. Rep-CH | Alt-EN vs. Rep-EN | Alt-CHEN vs. Rep-CHEN |
| df | 779 | 779 | 1599 |
| *t* value | 8.497 | 18.771 | 27.150 |
| *P* value | < 0.001 | < 0.001 | < 0.001 |
| Cohen's d | 0.304 | 0.672 | 0.679 |
| 95% CI | [0.056, 0.090] | [0.118, 0.146] | [0.133, 0.154] |
| | | **Experiment 4** | |
| Condition | Alt-CH vs. Rep-CH | Alt-EN vs. Rep-EN | Alt-CHEN vs. Rep-CHEN |
| df | 779 | 779 | 1599 |
| *t* value | 19.483 | 6.726 | 22.643 |
| *P* value | < 0.001 | < 0.001 | < 0.001 |
| Cohen's d | 0.698 | 0.241 | 0.566 |
| 95% CI | [0.102, 0.124] | [0.045, 0.082] | [0.096, 0.114] |
| | | **Experiment 5** | |
| Condition | Alt-CH vs. Rep-CH | Alt-EN vs. Rep-EN | Alt-CHEN vs. Rep-CHEN |
| df | 779 | 779 | 1599 |
| *t* value | 20.841 | 10.221 | 61.125 |
| *P* value | < 0.001 | < 0.001 | < 0.001 |
| Cohen's d | 0.746 | 0.366 | 1.528 |
| 95% CI | [0.120, 0.145] | [0.050, 0.073] | [0.211, 0.224] |
| | | **Experiment 6** | |
| Condition | Alt-KR vs. Rep-KR | Alt-IT vs. Rep-IT | Alt-KRIT vs. Rep-KRIT |
| df | 779 | 779 | 1599 |
| *t* value | 12.454 | 13.851 | 39.241 |
| *P* value | < 0.001 | < 0.001 | < 0.001 |
| Cohen's d | 0.446 | 0.496 | 0.981 |
| 95% CI | [0.082, 0.113] | [0.081, 0.108] | [0.179, 0.198] |
| | | **Experiment 7a and 8a** | |
| Condition | Alt-CH vs. Rep-CH | Alt-EN vs. Rep-EN | Alt-CHEN vs. Rep-CHEN |
| df | 779 | 779 | 1599 |
| *t* value | 11.327 | 8.549 | 29.376 |
| *P* value | < 0.001 | < 0.001 | < 0.001 |
| Cohen's d | 0.406 | 0.306 | 0.734 |
| 95% CI | [0.085, 0.120] | [0.066, 0.105] | [0.164, 0.188] |
| | | **Experiment 7a** | |
| Condition | Alt-CH vs. Rep-CH | Alt-EN vs. Rep-EN | Alt-CHEN vs. Rep-CHEN |
| df | 779 | 779 | 1599 |
| *t* value | 11.901 | 14.297 | 23.370 |
| *P* value | < 0.001 | < 0.001 | < 0.001 |
| Cohen's d | 0.426 | 0.512 | 0.584 |
| 95% CI | [0.096, 0.134] | [0.131, 0.173] | [0.138, 0.164] |
| | | **Experiment 8a** | |

| Condition | Alt-CH vs. Rep-CH | Alt-EN vs. Rep-EN | Alt-CHEN vs. Rep-CHEN |
|---|---|---|---|
| df | 779 | 779 | 1599 |
| *t* value | 12.791 | 4.655 | 20.260 |
| *P* value | < 0.001 | < 0.001 | < 0.001 |
| Cohen's d | 0.458 | 0.167 | 0.507 |
| 95% CI | [0.094, 0.129] | [0.028, 0.070] | [0.130, 0.158] |
| **Experiment 9** | | | |
| Condition | Alt-KR vs. Rep-KR | Alt-IT vs. Rep-IT | Alt-KRIT vs. Rep-KRIT |
| df | 779 | 779 | 1599 |
| *t* value | 10.206 | 3.853 | 14.925 |
| *P* value | < 0.001 | < 0.001 | < 0.001 |
| Cohen's d | 0.365 | 0.138 | 0.373 |
| 95% CI | [0.088, 0.130] | [0.020, 0.062] | [0.094, 0.122] |

*Notes.* CH, Chinese; EN, English; KR, Korean; IT, Italian.
